# A microplate screen to estimate metal-binding affinities of metalloproteins

**DOI:** 10.1101/820670

**Authors:** Patrick Diep, Radhakrishnan Mahadevan, Alexander F. Yakunin

## Abstract

Solute-binding proteins (SBPs) from ATP-binding cassette (ABC) transporters play crucial roles across all forms of life in transporting compounds against chemical gradients. Some SBPs have evolved to scavenge metal substrates from the environment with nanomolar and micromolar affinities (*K*_D_). There exist well established techniques like isothermal titration calorimetry for thoroughly studying these metalloprotein interactions with metal ions, but they are low-throughput. For protein libraries comprised of many metalloprotein homologues and mutants, and for collections of buffer conditions and potential ligands, the throughput of these techniques is paramount. In this study, we describe an improved method termed the microITFQ-LTA and validated it using *Cj*NikZ, a well-characterized nickel-specific SBP (Ni-BP) from *Campylobacter jejuni*. We then demonstrated how the microITFQ-LTA can be designed to screen through a small collection of buffers and ligands to elucidate the binding profile of a putative Ni-BP from *Clostridium carboxidivorans* that we call *Cc*SBPII. Through this study, we showed *Cc*SBPII can bind to various metal ions with *K*_D_ ranged over 3 orders of magnitude. In the presence of L-histidine, *Cc*SBPII could bind to Ni^2+^ over 2000-fold more tightly, which was 11.6-fold tighter than *Cj*NikZ given the same ligand.

**Highlights:**

- an improved version of a high-throughput screen (microITFQ-LTA) is described for multiplexed elucidation of metalloprotein binding profiles
- validation was accomplished with the previously characterized *Cj*NikZ; testing was accomplished with an uncharacterized homologue herein named *Cc*SBPII
- *Cc*SBPII is shown to bind to multiple transition metal ions with a large range of affinities, and potentially overcome mismetallation using a simple histidine metallophore

## Introduction

Metalloproteins are proteins whose function and structure involve interactions with metal ions. Several techniques in metal-binding studies can be used to study these interactions *in vitro*. The most widely accepted technique is isothermal titration calorimetry (ITC) as it can determine multiple thermodynamic parameters in a single binding experiment. However, obtaining meaningful ITC data is not a trivial undertaking. Requisites include an easily expressed protein in large quantities that is stable under prolonged stirring conditions (*viz*. it does not precipitate), the protein generates a heat signature upon binding that is appreciably larger than the heat of dilution, there is sufficient time to run multiple ITC trials to optimize the ratio of ligand and protein, and more. This last requirement can be especially cumbersome due to the time needed to equilibrate the system, but a value “c” can be calculated to better adjust this ratio if an accurate estimation of the protein’s binding constant *K*_D_ can be made using other techniques.^1,2^ Given that meaningful ITC data is obtained, one must consider whether ITC is the most appropriate technique to determine the binding constants for large protein families, mutant libraries, or both; and for multiple ligands and solvent conditions of interest. It is useful in such cases to consider a high-throughput screening technique, such as one that employs intrinsic tryptophan fluorescence quenching (ITFQ) in a ligand titration assay (LTA), herein termed ITFQ-LTA, to complement the lower throughput but more comprehensive ITC experiments.

Fluorescence quenching of a protein’s intrinsic fluorophores (*i.e*. tryptophan) upon sequential exposure to a ligand through step-wise titration can generate binding curves for estimating *K*_D_, which therefore complements ITC by helping to calculate c. To increase the throughput for more extensive library screening, microplate readers equipped with fluorescence-reading capabilities can be used instead of spectrofluorometers, which has been tested for interactions between albumins and proteases with organic compounds.^3–5^ Microplate-based ITFQ-LTA (microITFQ-LTA) has also been used to study interactions between metalloproteins and UO_2_^2+^, but these researchers’ decision to use Equation (**1**), the Stern-Volmer equation (variables described further below), to fit the fluorescence quenching data is questionable.^6,7^

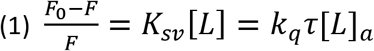

The Stern-Volmer equation models collisional quenching, a phenomenon where the excited-state fluorophore is returned to ground state through contact with an *external* quencher in a non-radiative manner. This typically occurs via spin-orbit coupling with paramagnetic entities.^8^ Proteinaceous tryptophans are often partially or fully buried in the hydrophobic core where these external quenchers have limited access, and cannot easily collide. Consequently, static quenching is an unlikely mechanism for protein fluorescence quenching too because it requires the external quencher to form a complex with the ground-state proteinaceous tryptophan.^9^ A stronger explanation is intramolecular self-quenching. Tryptophan’s large dipole moment causes it to be highly sensitive to changes in the polarity of its environment. Ligand-binding frequently induces conformational changes across the protein structure that changes the direction and magnitude of local electric fields around the tryptophans’ indole rings. This can lead to saturable quenching (and sometimes enhancement) of its fluorescence through increased non-radiative energy transfer to nearby moieties of the surrounding protein matrix, typically through electron transfer into the peptide backbone and nearby side chains.^9–14^

Regardless of the exact mechanism for protein fluorescence quenching, the association phenomenon (*i.e*. binding event) can be followed by the magnitude of the change in fluorescence in response to the ligand titrations, modelled by Equation (**2**) for 1:1 metal-protein complexes. This model uses basic mass-action kinetics that assumes the protein and ligand concentration that can freely form complexes at any point in time during the ITFQ-LTA is a sum of what is uncomplexed (*i.e*. free) and what is already complexed, described in Equations (**3**) and (**4**).^9,15,16^

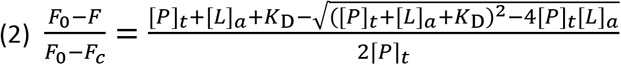

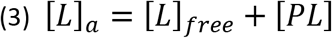

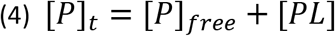

For all Equations in this study: *F*_0_ and *F*_c_ are the measured fluorescence when the protein is fully unsaturated and fully saturated, respectively. *F* is the measured fluorescence at a ligand titration step. *K*_sv_ is the Stern-Volmer quenching constant and *k*_q_ is the bimolecular quenching rate constant. τ is the fluorescence lifetime of the unquenched fluorophore. [*L*]_a_ and [*P*]_t_ are the concentrations of total ligand and protein, respectively. [*L*]_free_ and [*P*]_free_ are the concentrations of ligand and protein that are uncomplexed with each other, respectively. [*PL*] is the concentration of the protein complexed with the ligand, and *K*_D_ is the apparent dissociation constant under a specific buffering condition.

Depending on how strongly the protein’s tryptophan fluorophores respond to a binding event, which is based on the number of tryptophans in the protein and the extent of the ligand-induced conformational change, a large signal with low noise can be obtained for low micromolar protein concentrations. If the *K*_D_ for a ligand is below the protein concentration by at least a factor of 10, it is generally more appropriate to use Equation (**2**). If the *K*_D_ for a ligand is larger than the protein concentration, which means the range of ligand concentrations used in the microITFQ-LTA is in molar excess of protein concentration, [*PL*] can be neglected in Equation (**3**), which simplifies the mass-action kinetics underpinning Equation (**2**) to yield Equation (**5**).^9^

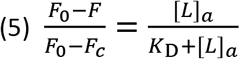

To simplify the variables for data analysis in this study, Equation (2) will be re-written as Equation (**6**), and Equation (**5**) will be re-written as Equation (**7**). *F*_obs_ is *F*_0_-*F* and *F*_max_ is *F*_0_-*F*_c_. When the Hill Coefficient “n” is used, [*L*]_a_ in both Equations (**6**) and (**7**) is substituted with [*L*]_a_^n^.

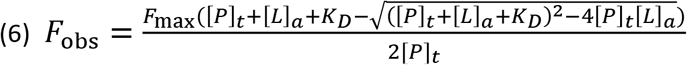

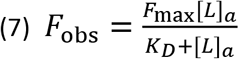

One group of metalloproteins are the solute-binding protein (SBP) components of metal ATP-binding cassette (ABC) transporters. These transporters are ubiquitous in nature as they provide cells the means to transport metal substrates against chemical gradients using the energy from nucleotide triphosphate hydrolysis. The specificity of ABC transporters is derived from its cognate SBP that contains the binding site for target metal ions.^17^ NikA from the *Escherichia coli* ABC transporter NikABCDE is the most thoroughly studied nickel-specific SBP (Ni-BP) with extensive biochemical and structural characterization.^18–26^ Other Ni-BPs have also been studied because of their implications in health, and include the pathogens *Campylobacter jejuni* (*Cj*NikZ)^27,28^, *Brucella suis* (*Bs*NikA)^28,29^, *Yersinia pestis* (*Yp*YntA)^28^, *Staphylococcus aureus* (*Sa*NikA, *Sa*CntA)^30^, *Helicobacter hepaticus* (*Hh*NikA)^31^, *Helicobacter pylori* (CeuE)^32^, *Vibrio parahaemolyticus* (*Vp*NikA)^33^, and *Clostridium difficile* (*Cd*OppA)^34^. However, it is not only pathogens that require metals to survive. *Clostridium carboxidivorans* is not known to be pathogenic, but possesses a cluster of coding sequences herein termed the acetogenesis/nickel acquisition (*ana*) operon that encodes a putative metal ABC transporter and acetogenesis-related enzymes that use metal co-factors (Supplementary Figure 1). This ABC transporter possesses two SBPs, herein termed *Cc*SBPI and *Cc*SBPII, which are both homologous to *Cj*NikZ (41% and 37%, respectively).

We are interested in studying and engineering Ni-BPs and their cognate ABC transporters for mining applications. In this study, we took a different approach from the aforementioned UO_2_^2+^-metalloprotein binding study and instead used Equations (**6**) and (**7**) to fit our microITFQ-LTA data. We first validated our method by closely replicating the *K*_D_ values for Ni^2+^➔*Cj*NikZ that have previously been determined using spectrophotometer-based ITFQ-LTA (specITFQ-LTA) and ITC. We then studied Ni^2+^➔*Cc*SBPII with different buffers and varied the ligands afterwards. *Cc*SBPII is an uncharacterized metalloprotein, but shares structural similarity to *Cj*NikZ (Supplementary Figure 2), so we narrowed our scope to elucidating its binding profile using the microITFQ-LTA screen so that it can be further studied with more comprehensive techniques like ITC and x-ray crystallography. Therefore, the focus of this study was to test and describe a microITFQ-LTA screening method for metalloproteins that builds on previous iterations of the method.^3–7^ We report *Cc*SBPII could bind to numerous transition metal ions with *K*_D_ values that ranged over 3 orders of magnitude, and could possibly deal with Cu^2+^ mismetallation by using L-histidine as a simple metallophore to selectively bind to the theoretical target Ni^2+^.

## Materials and Methods

### DNA synthesis, cloning, and strains

The open-reading frames encoding *Cc*SBPII from *Clostridium carboxidivorans* P7T (Ccar_RS11695, NCBI ID: NZ_CP011803.1) and *Cj*NikZ^28^ were synthesized (Twist Bioscience, San Francisco, USA) without start codons and without their native signal peptides identified using SignalP 5.0 with the Gram-positive setting.^35^ Their open-reading frames were cloned into the expression vector p15TV-L (AddGene ID: 26093) under the T7 promoter and in-frame with the N-terminal 6xHisTag (Twist Bioscience). Upon receiving the completed plasmids, they were transformed into the DH5α and LOBSTR (Kerafast, #EC1001) *Escherichia coli* strains, then plated onto LB-Agar (carbenicillin, 100 μg/mL). The plasmids were miniprepped (GeneAid, #PD300) into MilliQ water and sequence verified at the ACGT Sequencing Facility (Toronto, Canada). Glycerol stocks were prepared for storage at −80 °C.

### Protein expression and purification

Starter cultures for *Cj*NikZ and *Cc*SBPII were grown from glycerol stock in LB with carbenicillin (100 μg/mL) for 16 hrs overnight at 37 °C with shaking. Expression cultures were started by pre-warming TB media with 5% glycerol and carbenicillin (100 μg/mL) to 37 °C before 1% v/v inoculation with the starter culture, then grown for 3 hrs at 37 °C until addition of IPTG (BioShop, #IPT002) to 0.4 mM for induction. The expression cultures were then transferred to 16 °C and grown for 16 hrs overnight with shaking, then pelleted with centrifugation and transferred to conical vials for one freeze-thaw cycle at −20 °C.

Frozen cell pellets were thawed and resuspended in binding buffer (10 mM HEPES, 500 mM NaCl, 5 mM imidazole, pH 7.2) to a final volume of 100 mL, followed by addition of 0.25 g lysozyme (BioShop, #LYS702). Cell pellet mixtures were sonicated for 25 min (Q700 Sonicator, Qsonica) and clarified by centrifugation. The soluble layer (supernatant) was applied to a cobalt-charged (Co-)NTA resin (ThermoFisher, #88221) pre-equilibrated with binding buffer in a gravity-column set-up. Bound proteins were cleansed with wash buffer (10 mM HEPES, 500 mM NaCl, 25 mM imidazole, pH 7.2) and the bound protein was collected with elution buffer (10 mM HEPES, 500 mM NaCl, 250 mM imidazole, pH 7.2). Protein concentrations were determined by Bradford assay, and protein purity was determined by SDS-PAGE analysis and densitometry on Image Lab 6.0 (Bio-Rad).

Eluted protein was combined with 0.4 mg of in-house purified TEV protease and TCEP to 1 mM (BioShop, #TCE101), then transferred to a 10 kDa MWCO dialysis bag (ThermoFisher, #68100) for dialysis in 4 L dialysis buffer (10 mM HEPES, 1 mM TCEP, 1g/L Chelex 100, pH 7.2) at 4 °C with gentle stirring for 72 hrs. Dialyzed samples were then applied to a cobalt-charged NTA resin twice and concentrated to 2 mL by centrifugation using 30 kDa MWCO concentrators (Millipore Sigma, #GE28-9323-61). This concentrate was applied to a MonoQ 5/50 anion exchange resin (GE Healthcare, #17516601) using 10 mM HEPES, pH7.2 for the mobile phase, and 10 mM HEPES, 1M NaCl, pH 7.2 for elution. Eluted proteins were collected in 96 deep-well blocks and stored at 4 °C during SDS-PAGE analysis. Selected fractions with sufficient purity were transferred to a 10 kDa MWCO dialysis bag for dialysis in 4 L dialysis buffer (10 mM HEPES, pH 7.2) at 4 °C with gentle stirring for 24 hrs and transferred to fresh dialysis buffer for another 24 hrs. Finally, they were flash-frozen drop-wise in liquid nitrogen before storage at −80 °C. Prior to usage in preliminary microITFQ-LTAs, the proteins were checked using circular dichroism, MALDI-ToF, and ICP-MS to ensure proper folding, molecular weight (*i.e*. successful 6x HisTag removal), and minimal metal contamination, respectively (Supplementary Information, Appendix A; Supplementary Figures 3 & 4).

### Preparation of activity buffers and ligand solution

Except for *Cj*NikZ which used either 20mM MOPS pH 8 or 40 mM HEPES pH 7.5 as the activity buffer to mimic conditions used in prior studies^27,28^, *Cc*SBPII used 10 mM HEPES pH 7.2 as the default activity buffer with adjustments made as prescribed by the type of microITFQ-lTA (*i.e*. HEPES concentration range, pH range, NaCl range). Activity buffers were adjusted to the desired pH using NaOH and HCl, then filter sterilized by syringe using a 0.2 μm PTFE membrane disc (PALL ID: 4187). Ligand solutions were created using the appropriate activity buffer as prescribed by the type of microITFQ-LTA to match the activity buffer in the well that the protein was diluted into (*e.g*. 1 mM NiCl_2_ was prepared in 10 mM HEPES pH 7.2 from a 1M stock solution of NiCl_2_ dissolved in de-ionized water, in order to be added to *Cc*SBPII diluted into 10 mM HEPES pH 7.2). Metal salts with >98% purity were used to prepare all ligand solutions requiring NiCl_2_, NiSO_4_, CoCl_2_, CuCl_2_, ZnCl_2_, MnCl_2_, and CdCl_2_. Ni-His_2_, Co-His_2_, Cu-His_2_ were each prepared with a 1:2 stoichiometric ratio using the metal chloride salt and L-histidine. Ni-Imi_2_, Ni-Gly_2_, and Ni-Imi_2_Gly_2_ were prepared with a 1:2, 1:2, and 1:2:2 stoichiometric ratio using NiCl_2_, imidazole (imi), and glycine (gly). Ni_2_-Citrate_2_, Ni-EDTA, and Ni-DETA_2_ were prepared with a 2:2, 1:1, and 1:2 stoichiometric ratio. All ligand solutions were prepared fresh and immediately used in microITFQ-LTAs; activity buffers were stored at 4 °C.

### Microplate-based intrinsic tryptophan fluorescence quenching – ligand titration assay (microITFQ-LTA)

microITFQ-LTAs were performed in black, opaque 96-well microplates (Greiner Bio-One, #655076) using the SpectraMax M2 spectrophotometer (Molecular Devices) with the following settings: top-read, 25 °C, λ_ex_ 280 nm, λ_em_ 380 nm, PMT high, sensitivity 30, carriage speed high, 5 s shaking before endpoint reads, and 3 s shaking in between reads during kinetic reads. *Cj*NikZ and *Cc*SBPII were combined with the appropriate activity buffer as prescribed by the specific microITFQ-LTA to 0.4 μM for a total volume of 200 μL and allowed to equilibrate with intermittent shaking for up to 15 min, or until the signal stabilized. After equilibration, aliquots of the ligands are added in stepwise fashion using a multichannel pipette up to a maximum of 120 μL when saturation was not reached, otherwise less. Mixing by pipette was avoided to prevent the introduction of air bubbles or the retaining of droplets inside the tips that would distort the signal. Therefore, each titration was followed by a 30 s equilibration with intermittent shaking before endpoint reads were made to ensure proper mixing. All trials were done in triplicates. To account for possible dilution effects in every trial, we included a triplicate negative control where the protein was titrated with only activity buffer (no metal ligand) using the same volumes added at each titration. A minor IFE was exhibited by histidine, which was accounted for by subtracting its individual fluorescence signal at λ_em_ 380 nm (obtained by titrating histidine in the absence of metals or protein) from each titration step. Otherwise, all ligand solutions and components of activity buffers did not exhibit the IFE for the concentrations used.

### Data analysis and visualization

All raw data is prepared in Microsoft 365 Excel for further analysis. Triplicate data for tests and negative controls (as described above) were averaged for each titration step (error propagation applied). The averaged negative control data was then subtracted from the averaged ligand test data at each titration step to remove the background signal and obtain *F*. The value of *F* before any titration is made, where the protein is unsaturated, is *F*_0_. To obtain a *F*_obs_ value for each titration step, *F* from all titration steps are subtracted from *F*_0_ such that the new *F*_0_ is zero and an up-right curve can be generated. These sets of x/y data pairs (*e.g*. [*L*]_a_/F_obs_, respectively) are inputted into GraphPad Prism 5.0 for determination of best-fit parameter values and confidence intervals through non-linear least-squares (NLLS) regression using Equation (**6**). This is performed with and without the hill coefficient n as needed, where [*L*]_a_ is substituted with [*L*]_a_^n^. The Kemmer and Keller (KK) method, described elswhere^36^, is used in Excel to compute an NLLS regression using Equation (**6**) or (**7**) for determining best-fit parameter values and 95% confidence intervals, including n. In all NLLS regressions, the minimization parameter is χ^2^, in which the residuals are weighted by observed variability of *F*_obs_ (1/σ^2^). All figures are produced using GraphPad Prism 5.0 (or for brevity, “Prism”).

## Results and Discussion

### microITFQ-LTA closely replicates published binding constants for CjNikZ and supports crystal structure data

The Ni-BP *Cj*NikZ belongs to NikZYXWV, a nickel ABC transporter, and has published *K*_D_ values of 0.61 ± 0.07 μM and 1.6 ± 0.2 μM (89% difference), determined by a specITFQ-LTA^27^ and by ITC^28^, respectively. In these studies, non-linear least squares (NLLS) regressions were performed to obtain these best-fit parameter values with the commercial software packages SigmaPlot and MicroCal Origin. To fit the data, one-binding site models (n = 1) were used. In this study, we determined the *K*_D_ value of *Cj*NikZ in the same buffers used in these studies: 20 mM MOPS pH 8, and 40 mM HEPES pH 7.5. We also used the same nickel salt to prepare the ligand solutions: NiCl_2_ and NiSO_4_, respectively.

Assuming *Cj*NikZ has one binding site (n = 1), and given that the concentration of nickel to be titrated is in molar excess of the protein concentration used (0.4 μM), Equation (**7**) was chosen to model the data using Prism (Figure 1). In 20 mM MOPS pH 8, we find that Ni^2+^➔*Cj*NikZ yields a *K*_D_ value of 2.23 μM with a confidence interval (CI) of 1.53 to 2.92 μM. In 40 mM HEPES pH 7.5, we find that Ni^2+^➔*Cj*NikZ yields a *K*_D_ value of 3.34 μM with a CI of 2.41 to 4.26 μM. This difference in *K*_D_ values matches the difference in the published *K*_D_ in that the MOPS condition had a slightly lower *K*_D_ than the HEPES condition. Since crystallographic data shows that *Cj*NikZ has only one binding site for Ni^2+^, we tested our first assumption here and included n in Equation (**7**) for a separate analysis. We found n to be 0.90 with a CI of 0.60 to 1.21, and similar *K*_D_ and *F*_max_ values, suggesting that the assumption is correct. Given that *Cj*NikZ’s *K*_D_ value of 1.6 μM determined by ITC is the most accurate value, the observed *K*_D_ values of 2.23 μM and 3.34 μM are only 33% and 70% different, respectively, which suggests the microITFQ-LTA is as accurate as the specITFQ-LTA that yielded an 89% difference.

**Figure 1.**
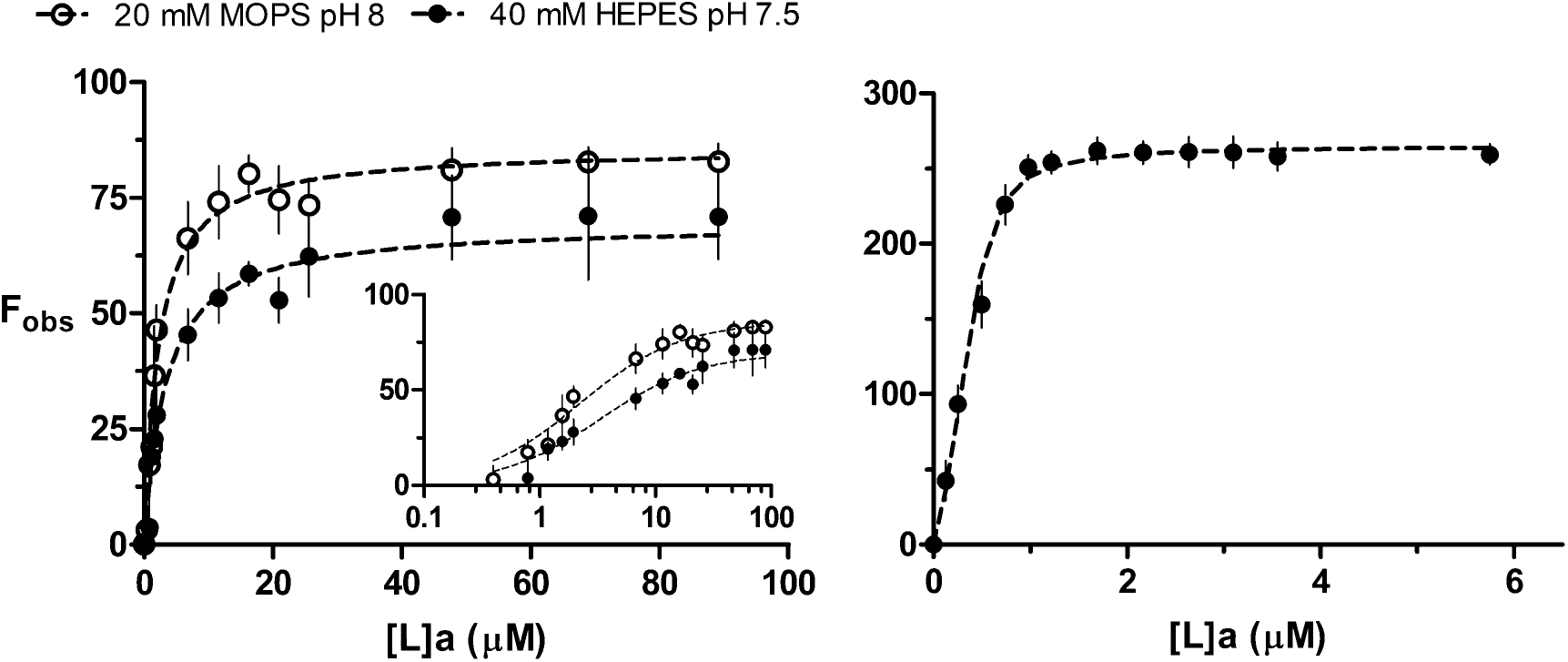
Binding curve for a Ni^2+^➔*Cj*NikZ titration (left-panel) in 20 mM MOPS pH 8 (empty circles) and 40 mM HEPES pH 7.5 (filled circles), as well as a (1:2)Ni-His➔*Cj*NikZ titration (right-panel) in 40 mM HEPES pH 7.5. Curves of best-fit are generated with Equation (**7**) on Prism (left-panel), and the KK method with Equation (**6**) and n on Excel (right-panel). An x-axis logarithmic plot of Ni^2+^➔*Cj*NikZ is provided for closer visual inspection (left-panel inset).

While Prism can model data with Equation (**7**), it cannot be used with Equation (**6**) for scenarios where the *K*_D_ for a ligand is below the protein concentration in the assay because this requires the inclusion of a second independent variable [*P*]_t_ that changes at each titration due to dilution. Therefore, we utilized a method described by Kemmer & Keller (2010) that combines Excel’s Solver function for NLLS regression with a simple plug-and-chug routine using Fisher’s *F* distribution to determine the CIs for the best-fit parameter values, herein termed the KK method for brevity. Using Equation (**7**) with the data for Ni^2+^➔*Cj*NikZ in 40 mM HEPES pH 7.5, we found Prism and Solver’s NLLS regression algorithms generated nearly identical best-fit parameters values (Table 1). Comparing the CIs computed by Prism and the KK method, we found they were also nearly identical, suggesting the KK method was a reliable protocol for modelling data with Equation (**7**) on Excel. Applying the KK method with Equation (**6**) instead for this data, we find there is little difference made to the best-fit parameter values, indicating that [*P*]_t_ was indeed negligible for characterizing this particular binding interaction between Ni^2+^ and *Cj*NikZ, thus confirming the assumption made earlier to choose Equation (**7**).

**Table 1.** Binding parameters for *Cj*NiKZ

| **Ni → *CjNikZ*, 40 mM HEPES pH 7.5** |  |  |  |  |  |  |  |  |  |  |  |
| --- | --- | --- | --- | --- | --- | --- | --- | --- | --- | --- | --- |
| NLLS Method | Model | $\chi^2$ | $n$ | n, 95% CI | | $K_D$ (μM) | $K_D$, 95% CI | | $F_{max}$ | $F_{max}$, 95% CI | |
|  |  |  |  | Lower | Upper |  | Lower | Upper |  | Lower | Upper |
| Prism | Eq.(7) | 4.45 | 1 |  |  | 3.34 | 2.41 | 4.26 | 69.40 | 64.64 | 74.17 |
|  |  | 4.32 | 0.90 | 0.60 | 1.21 | 3.39 | 2.39 | 4.39 | 72.95 | 58.93 | 86.96 |
| KK | Eq.(7) | 4.45 | 1 |  |  | 3.34 | 2.57 | 4.27 | 69.41 | 65.32 | 74.14 |
|  |  | 4.31 | 0.91 | 0.67 | 1.26 | 3.39 | 2.63 | 4.41 | 72.94 | Error<sup>a</sup> |  |
| KK | Eq.(6) | 4.56 | 1 |  |  | 3.02 | 2.24 | 3.92 | 68.59 | 64.49 | 73.37 |
|  |  | 4.32 | 0.88 | 0.65 | 1.23 | 3.08 | 2.34 | 4.10 | 73.03 | Error<sup>a</sup> |  |

| **(1:2)<sup>b</sup> Ni-His → *CjNikZ*, 40 mM HEPES pH 7.5** |  |  |  |  |  |  |  |  |  |  |  |
| --- | --- | --- | --- | --- | --- | --- | --- | --- | --- | --- | --- |
| NLLS Method | Model | $\chi^2$ | $n$ | n, 95% CI | | $K_D$ (μM) | $K_D$, 95% CI | | $F_{max}$ | $F_{max}$, 95% CI | |
|  |  |  |  | Lower | Upper |  | Lower | Upper |  | Lower | Upper |
| Prism | Eq.(7) | 53.17 | 1<sup>a</sup> |  |  | 0.30 | 0.13 | 0.47 | 293 | 265.20 | 320.80 |
|  |  | 6.74 | 2.17 | 1.71 | 2.63 | 0.090 | 0.038 | 0.14 | 264.7 | 256.90 | 272.50 |
| KK | Eq.(7) | 53.17 | 1<sup>a</sup> |  |  | 0.30 | 0.19 | 0.45 | 293.02 | 267.65 | 328.05 |
|  |  | 6.74 | 2.17 | 1.83 | 2.58 | 0.090 | 0.056 | 0.13 | 264.69 | 258.70 | 271.24 |
| KK | Eq.(6) | 16.38 | 1<sup>a</sup> |  |  | 0.093 | 0.050 | 0.15 | 272.36 | 261.22 | 284.48 |
|  |  | 4.42 | 1.32 | 1.11 | 1.57 | 0.051 | 0.031 | 0.074 | 265.25 | 255.23 | 278.65 |

a. CI could not be computed due to an inability to define a local minimum (refer to Supplementary Figures 5)
b. a 1:2 stoichiometric ligand solution of NiSO<sub>4</sub> and L-histidine

Interestingly, CIs could not be computed by the KK method for Equations (**6**) and (**7**) when n was included. Overparameterization can be diagnosed through the KK method using its χ^2^ plots. Here, a value χ^2^_bf_ is calculated using Fisher’s F distribution and the bounds for the 95% CI of a best-fit parameter value are defined by where the curves have a y-value equal to χ^2^_bf_, based on a polynomial function. Therefore, a requisite for the KK method to be used is for the slope of the curve on both sides to be sufficiently steep to define a reasonable CI. We generated χ^2^ plots against *K*_D_ and *F*_max_ (Supplementary Figure 5) for both models and found the inclusion of n reduced χ^2^ by 4-6% (*viz*. marginally improved the goodness of fit), but at the expense flattening the curve for *F*_max_ to the point where a reasonable CI could not be calculated, which is symptomatic of overparameterization of the model by n.^36^ Overall, this indicates n should be used judiciously, and only when minimizations of χ^2^ are substantial and biologically meaningful.

In Lebrette *et al*. (2014), they complement their ITC data for *Cj*NikZ with three crystal structures, one of which includes *Cj*NikZ bound to Ni^2+^ chelated by a histidine (PDB ID: 4OEU). Upon closer analysis, a (very) well-formed octahedral coordination complex is created between *Cj*NikZ, Ni^2+^, and histidine. However, their ITC data revealed that *Cj*NikZ did not produce any thermic event when Ni(His)_2_ was injected. While they explain the histidine in the crystal structure was an artifact of their crystallization procedure, we performed a Ni(His)_2_➔*Cj*NikZ microITFQ-LTA in 40 mM HEPES pH 7.5, the same buffer conditions for the ITC they performed, and detected a low nanomolar binding event. Using the KK method with Equation (**6**) and inclusion of n, we observed a strong binding event: n of 1.32 with a CI of 1.11 to 1.57, *K*_D_ of 51 nM (!) with a CI of 31 to 74 nM, and an *F*_max_ of 265.25 with a CI of 255.23 to 278.65 (Table 1). Of pertinence, we subtracted the background fluorescence emission of histidine at each titration step through an additional set of controls to remove the IFE. Nonetheless, the IFE contribution of histidine only began appearing in the last two data points reported in Figure 1. We were not surprised by this finding because Ni-BPs have been found to bind to Ni-His complexes before with nanomolar affinity, which includes other structures solved in the same study by Lebrette *et al*. (2014). It is possible that in their study a small amount of Ni(His)_2_ was accidentally exposed to *Cj*NikZ in the sample cell upon insertion of the syringe into the sample, or by some other means, which would have saturated the protein prior to the first injection.

Compared to the usage of n for Ni^2+^➔*Cj*NikZ in 40 mM HEPES pH 7.5 where it had a detrimental role on computing CIs by the KK method, n plays an important role in properly fitting Equations (**6**) and (**7**) to the data here. This is illustrated in Supplementary Figure 6 where the curves for both *K*_D_ and *F*_max_ were lowered and slightly narrowed by unconstraining n (*viz*. including it in the model), and Supplementary Figure 7 where the curve for n was lowered and noticeably narrowed when applied to Equation (**6**). By including n, χ^2^ was reduced by 88% and 73% for Equation (**7**) and (**6**), respectively. Overall, this provides evidence supporting the functional role of the histidine in assisting *Cj*NikZ in binding Ni^2+^ more tightly and helps explain why histidine is found inside the crystal structure of *Cj*NikZ. Next, we demonstrate the microITFQ-LTA’s utility in understanding how *Cc*SBPII binds to various metal ligands under different conditions. The best-fit parameter values are summarized in Table 2.

**Table 2.** Binding parameters for *Cc*SBPII

| Buffer <sup>a</sup> | n | n, 95% CI | | $K_D$<br>(μM) | $K_D$, 95% CI | | $F_{max}$ | $F_{max}$, 95% CI | |
| --- | --- | --- | --- | --- | --- | --- | --- | --- | --- |
|  |  | Lower | Upper |  | Lower | Upper |  | Lower | Upper |
| 10 mM HEPES pH 7.2 |  |  |  | 13.40 | 10.71 | 16.09 | 178.42 | 170.22 | 186.63 |
| 10 mM MOPS pH 7.2 |  |  |  | 25.83 | 19.68 | 31.98 | 196.89 | 185.51 | 208.27 |
| 10 mM PIPES pH 7.2 |  |  |  | 22.08 | 17.66 | 26.50 | 185.58 | 175.99 | 195.17 |
| 10 mM Tris pH 7.2 |  |  |  | 16.18 | 13.90 | 18.47 | 169.33 | 163.05 | 175.60 |
| 10 mM Phosphate pH 7.2 |  |  |  |  |  | Error <sup>b-1</sup> |  |  |  |
| 10 mM HEPES pH 7.2 |  |  |  | 12.67 | 10.77 | 14.56 | 174.90 | 167.61 | 182.18 |
| 20 mM HEPES pH 7.2 |  |  |  | 12.24 | 10.19 | 14.29 | 168.05 | 163.10 | 172.99 |
| 40 mM HEPES pH 7.2 |  |  |  | 15.85 | 12.74 | 18.96 | 160.09 | 152.90 | 167.28 |
| 100 mM HEPES pH 7.2 |  |  |  | 30.76 | 22.78 | 38.75 | 152.81 | 142.81 | 162.80 |
| 1000 mM HEPES pH 7.2 |  |  |  | 19.36 | 4.81 | 33.92 | 23.38 | 18.57 | 28.18 |
| 10 mM HEPES 0 mM NaCl, pH 7.2 |  |  |  | 13.75 | 11.52 | 15.98 | 185.31 | 177.85 | 192.77 |
| 10 mM HEPES 10 mM NaCl, pH 7.2 |  |  |  | 18.47 | 15.03 | 21.92 | 181.64 | 173.42 | 189.86 |
| 10 mM HEPES 100 mM NaCl, pH 7.2 |  |  |  | 46.65 | 37.60 | 55.70 | 172.18 | 161.62 | 182.75 |
| 10 mM HEPES 1000 mM NaCl, pH 7.2 |  |  |  | 77.35 | 34.15 | 120.54 | 143.16 | 96.54 | 189.78 |
| 10 mM HEPES pH 2.5 |  |  |  |  |  | Error <sup>b-2</sup> |  |  |  |
| 10 mM HEPES pH 6.5 |  |  |  | 39.24 | 32.68 | 45.79 | 142.87 | 136.65 | 149.08 |
| 10 mM HEPES pH 7.0 |  |  |  | 22.34 | 18.63 | 26.04 | 155.04 | 147.61 | 162.48 |
| 10 mM HEPES pH 7.2 |  |  |  | 13.76 | 11.59 | 15.92 | 175.61 | 167.44 | 183.77 |
| 10 mM HEPES pH 7.5 |  |  |  | 12.31 | 10.20 | 14.42 | 171.84 | 165.46 | 178.22 |
| 10 mM HEPES pH 8.0 |  |  |  | 5.39 | 3.89 | 6.88 | 204.48 | 196.83 | 212.12 |

| Ligand <sup>c</sup> | n | n, 95% CI | | $K_D$<br>(μM) | $K_D$, 95% CI | | $F_{max}$ | $F_{max}$, 95% CI | |
| --- | --- | --- | --- | --- | --- | --- | --- | --- | --- |
|  |  | Lower | Upper |  | Lower | Upper |  | Lower | Upper |
| NiSO<sub>4</sub> |  |  |  | 14.13 | 13.27 | 14.98 | 182.51 | 178.74 | 186.28 |
| NiCl<sub>2</sub> |  |  |  | 12.00 | 11.12 | 12.88 | 176.68 | 172.70 | 180.66 |
| CoCl<sub>2</sub> |  |  |  | 21.11 | 13.62 | 28.60 | 95.99 | 86.11 | 105.87 |
| CuCl<sub>2</sub> |  |  |  | 8.54 | 5.81 | 11.28 | 595.10 | 552.82 | 637.38 |
| MnCl<sub>2</sub> |  |  |  | 98.00 | 83.00 | 113.01 | 104.10 | 98.39 | 109.80 |
| ZnCl<sub>2</sub> |  |  |  | 153.89 | 85.88 | 221.90 | 236.20 | 211.06 | 261.35 |
| CdCl<sub>2</sub> |  |  |  | 11.61<sup>d</sup> | 8.70 | 14.52 | 259.93 | 240.40 | 279.47 |
| (1:2) Ni-His <sup>f</sup> | 0.83 | 0.80 | 0.86 | 4.40<sup>e</sup> | 1.33 | 8.76 | 350.42 | 347.22 | 354.06 |
| (1:2) Co-His <sup>f</sup> | 1.21 | 1.13 | 1.29 | 557.85<sup>e</sup> | 509.88 | 608.20 | 439.38 | 430.69 | 448.83 |
| (1:2) Cu-His <sup>f</sup> | 1.19 | 1.02 | 1.38 | 36.08<sup>e</sup> | 14.69 | 65.45 | 425.31 | 414.08 | 437.36 |
| (1:2) Ni-Imi <sup>f</sup> | 0.82 | 0.77 | 0.88 | 6.76 | 6.00 | 7.51 | 222.32 | 216.74 | 227.90 |
| (1:2) Ni-Gly <sup>f</sup> | 0.80 | 0.70 | 0.90 | 13.00 | 9.89 | 16.10 | 348.06 | 335.77 | 360.35 |
| (1:2:2) Ni-Imi-Gly <sup>f</sup> | 0.86 | 0.74 | 0.98 | 14.35 | 10.37 | 18.32 | 367.10 | 343.56 | 390.64 |

Best-fit parameter values and 95% confidence intervals (CI) are by default calculated on Prism without inclusion of n, meaning that n = 1, and using Equation (7) unless otherwise indicated.
a. ligand used is Ni<sup>2+</sup> prepared with NiCl<sub>2</sub>
b-1. saturation was not achieved in the ligand range titrated
b-2. no change in fluorescence occurred compared to the controls
c. 10 mM HEPES pH 7.2 is used in all cases for CcSBPII and to prepare the ligand solutions
d. values and corresponding lower and upper bound for CI is reported in millimolar (mM)
e. values and corresponding lower and upper bound for CI is reported in nanomolar (nM)
f. the KK method with inclusion of n, meaning n is unconstrained, is used with Equation (6) instead
His: L-histidine, Gly: L-glycine, Imi: imidazole

### microITFQ-LTAs for buffer screening with CcSBPII and nickel

It is well known that pH buffers are crucial for numerous biochemical, cellular, and environmental assays; it is also known that buffer molecules can interact with the sample and confound the analysis, which is especially true for metal-binding studies. We performed (NiCl_2_) Ni^2+^➔*Cc*SBPII microITFQ-LTAs with various Good’s buffers claimed to be “metal-compatible” buffers^37^, as well as phosphate buffer (Figure 2). Using Equation (**7**) in Prism with n = 1, 10 mM HEPES, PIPES, MOPS, and Tris at pH 7.2 all generated similar binding curves with *K*_D_ values ranging from 13 to 25 μM (Table 2). These buffers all possess lone pair electron orbitals at pH 7.2 and can therefore act as Lewis bases to form coordinate bonds with Ni^2+^.^38^ While this may explain the minor differences in *K*_D_ between the four buffers, their CIs show some overlap. This is also true for the *F*_max_ values and their CIs. It is likely that the microITFQ-LTAs could not distinguish between the effects the four buffers had on Ni^2+^➔*Cc*SBPII, which indicated to us that the use of any of these four buffers is appropriate since saturation could be achieved, as long the selected buffer is consistently used in further assays. A binding curve could not be generated with the phosphate buffered likely because metal-phosphate complexes tend to precipitate, thus preventing *Cc*SBPII from interacting with the metal.^39^ Therefore, we continued testing HEPES as the buffer of choice because it provided the lowest *K*_D_, suggesting the buffer interfered the least with the Ni^2+^➔*Cc*SBPII interaction.

**Figure 2.**
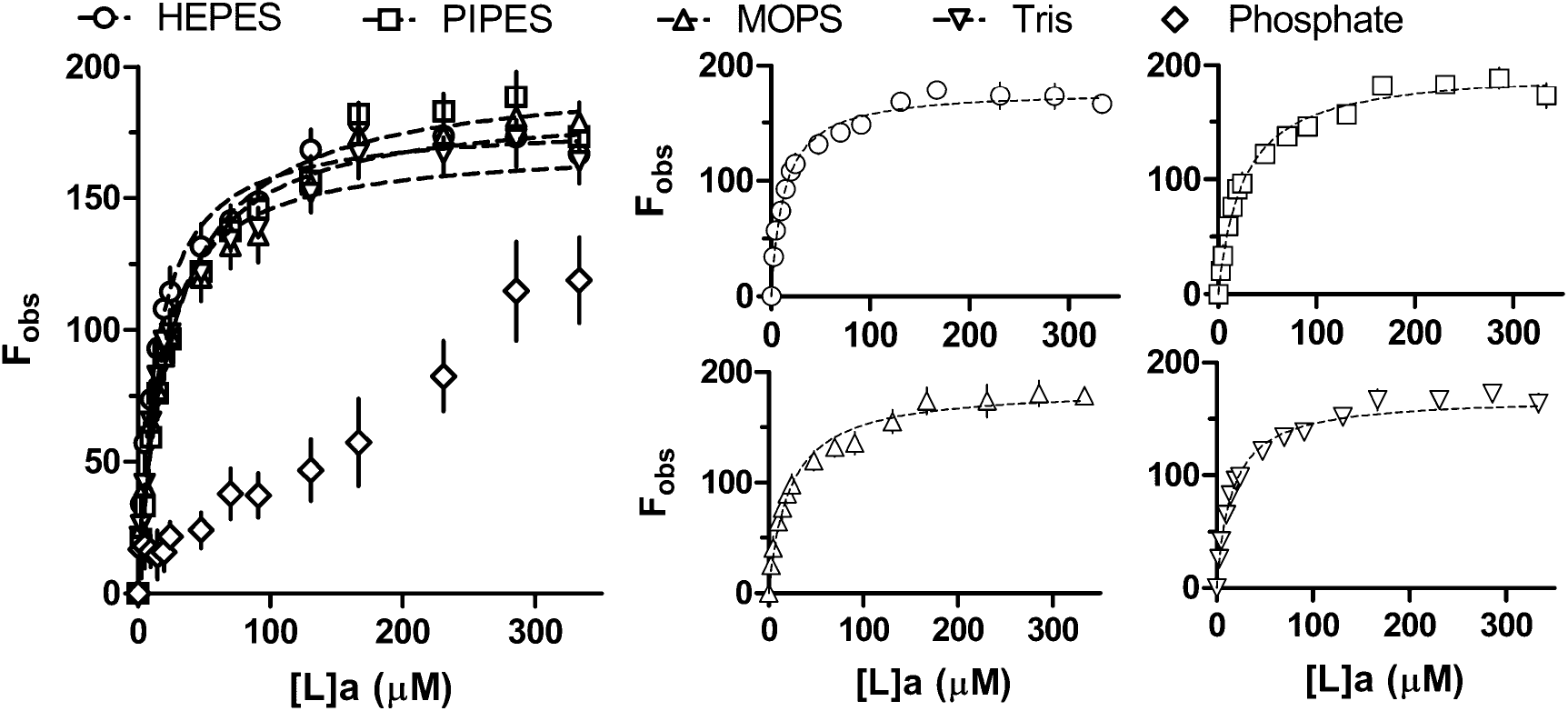
Binding curves for Ni^2+^➔*Cc*SBPII titrations (left-panel) at pH 7.2 for 10 mM HEPES (circle), 10 mM PIPES (square), 10 mM MOPS (upward triangle), 10 mM Tris (downward triangle), and 10 mM phosphate (diamond). Right panels show the same curves in the left panel individually (except 10 mM phosphate) for easier visual inspection.

When varying [HEPES] from 10 – 1000 mM at pH 7.2 and using Equation (**7**) in Prism with n = 1, we found the binding curves changed in two ways (Figure 3). *K*_D_ increased up to 2.4-fold over a change in 1-2 orders of magnitude of [HEPES], possibly because the higher concentrations of HEPES molecules effectively siphoned Ni^2+^ from the pool available for *Cc*SBPII to bind to, meaning that [*L*]_free_ for each titration step was overestimated and lead to a rightward-positive shift of the binding curve. More notably, *F*_max_ decreased 7.5-fold over a change in 2 orders of magnitude of [HEPES]. Specifically *F*_0_, which is needed to calculate *F*_max_, decreased and indicates the fluorescence from tryptophans had already been reduced prior to the first titration, noting here that 1000 mM HEPES displayed a negligible increase in absorbance at the wavelengths used compared to 10 mM HEPES (data not shown). At higher concentrations, it is likely that HEPES exerts a chaotropic effect on the *Cc*SBPII’s hydration shell, thus partially denaturing *Cc*SBPII. This could alter the environments surrounding the tryptophans as well as the binding cavity, which could also explain why *K*_D_ appeared to increase.^40,41^ Circular dichroism, thermal shift assays, or any method to probe changes in the secondary structure are needed to explore this. However, [HEPES] in the 1000 mM range is not typical of or practical for biochemical experiments and it was simpler to choose a lower [HEPES] with sufficient pH buffering capacity for the microITFQ-LTAs, so we continued with 10 mM HEPES.

**Figure 3.**
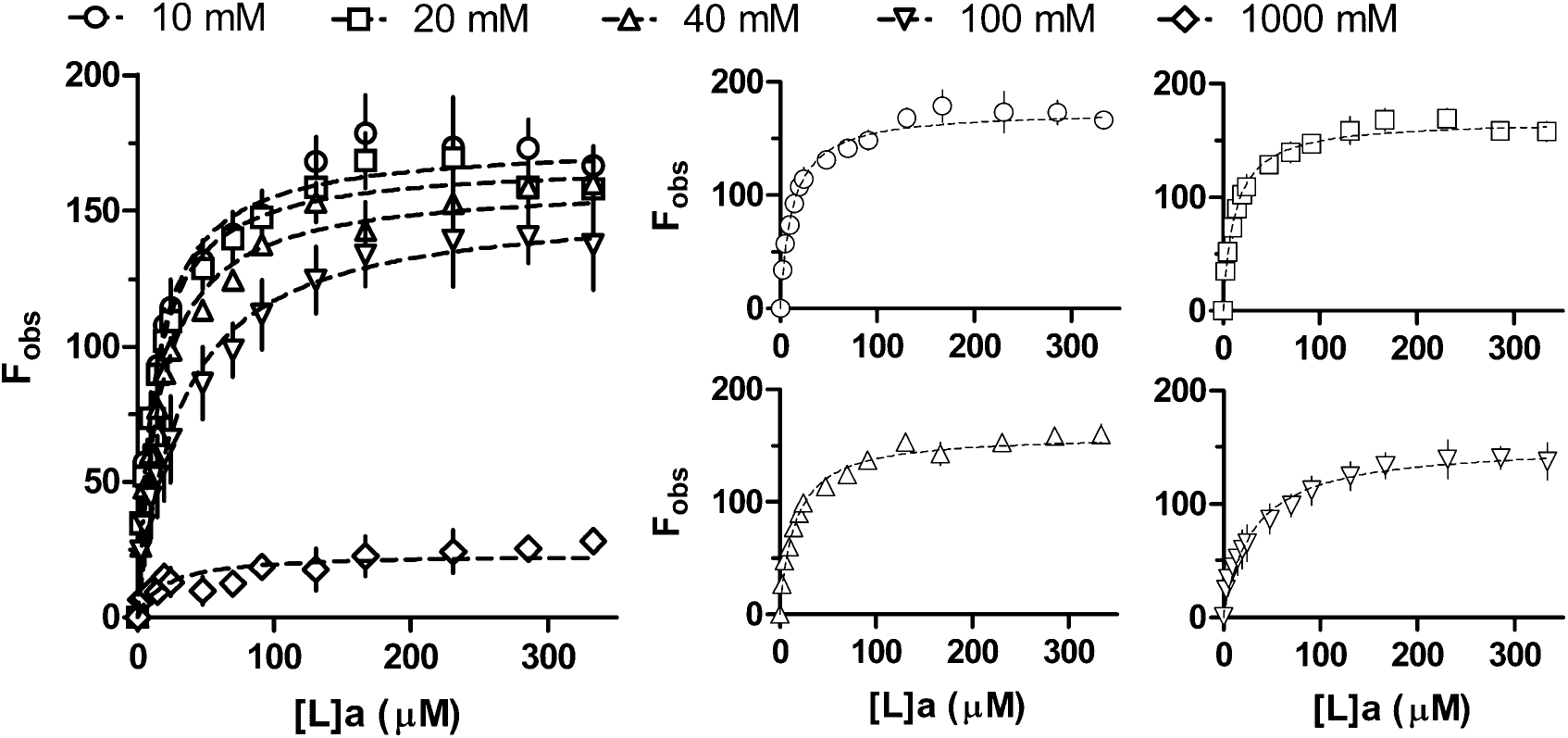
Binding curves for Ni^2+^➔*Cc*SBPII titrations (left-panel) at pH 7.2 with HEPES at 10 mM (circle), 20 mM (square), 40 mM (upward triangle), 100 mM (downward triangle), and 1000 mM (diamond). Right panels show the same curves in the left panel individually (except 1000 mM HEPES) for easier visual inspection.

When varying [NaCl] from 0 to 1000 mM at pH 7.2, and using Equation (**7**) in Prism with n = 1, we found the binding curves changed in a similar manner to when [HEPES] was varied (Figure 4). *K*_D_ increased 5.6-fold and *F*_max_ decreased 1.3-fold from 0 to 1000 mM NaCl, which likely reflects chaotropic effects on *Cc*SBPII’s hydration shell exerted by high [NaCl]. *C. carboxidvorans* is not known to be a halophile, meaning that its proteins are unlikely to form salt-bridge networks on their surfaces that would otherwise stabilize the structure. The more pronounced effect on *K*_D_ compared to the case with high [HEPES] may be due to Na^+^. While it is a hard ion that unlikely competes directly with Ni^2+^ to form coordinate bonds with *Cc*SBPII’s binding residues, its positive charge could still lead to electrostatic interactions with the negatively-charged binding cavity (Supplementary Figure 8), thus hindering the entrance and binding of Ni^2+^. Therefore, we chose not to include [NaCl] to minimize the number of buffer components that could complicate the Ni^2+^➔*Cc*SBPII interaction and continued with 10 mM HEPES.

**Figure 4.**
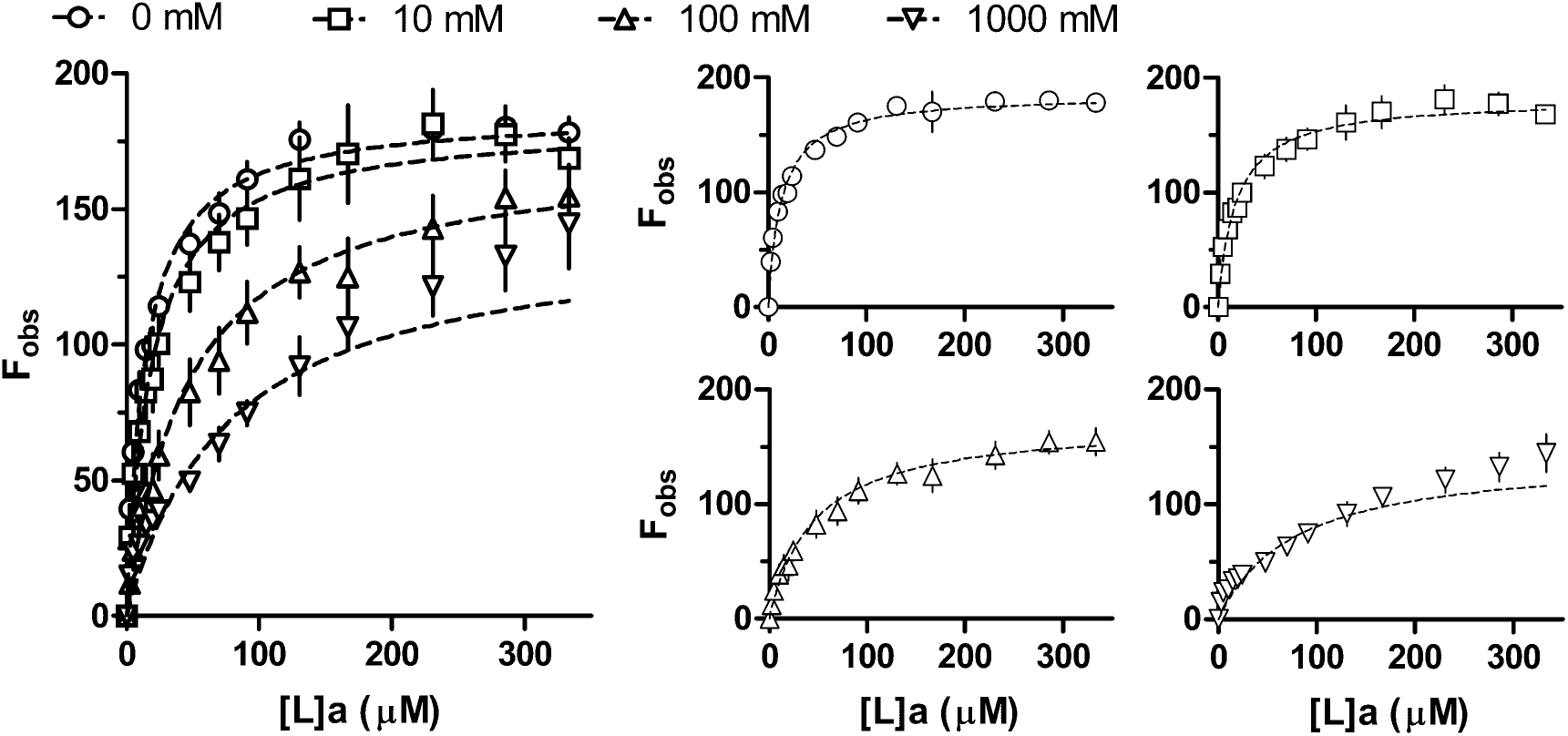
Binding curves for Ni^2+^➔*Cc*SBPII titrations (left-panel) at pH 7.2 with 10 mM HEPES and NaCl at 0mM (circle), 10 mM (square), 100 mM (upward triangle), and 1000 mM (downward triangle). Right panels show the same curves in the left panel individually (except pH 2.5 and pH 8.0) for easier visual inspection.

When varying the pH from 6.5 to 8.0 for 10 mM HEPES, and using Equation (**7**) in Prism with n = 1, we found *K*_D_ decreased 7.3-fold and *F*_max_ increased 1.4-fold in contrast to increasing [HEPES] and [NaCl], indicating a stronger binding event in more alkaline conditions (Figure 5). No change in fluorescence was observed at pH 2.5. Generally, pH plays a crucial role in metalloprotein function because it affects the ionization of the sidechains involved in binding to metal ions.^40,42^ Two of the four hypothesized binding residues based on structural superimposition with *Cj*NikZ (Supplementary Figure 2) are histidines that have a pK_a_ of 6.0 for the imidazole side chain; a pH below this value would protonate their nitrogens rendering them unable to contribute the lone pairs of electrons to form coordinate bonds with Ni^2^. With increasing pH from 6.5 to 8.0, these histidines could become increasingly deprotonated such that coordinate bonds can be formed more easily with Ni^2+^. The other two hypothesized binding residues are lysine (pK_a_ 10.54) and arginine (pK_a_ 12.48), which may experience some deprotonation in the pH range tested that would allow for better coordinate bond formation, though this effect is likely minor due to their large pK_a_ values. To avoid *Cc*SBPII’s isoelectric point (pI 5.78) and to limit nickel hydroxide precipitation beginning at pH 8.5 to 9,^43^ we defined 10 mM HEPES at pH 7.2 as a suitable buffer condition to conduct further ligand screening for *Cc*SBPII.

**Figure 5.**
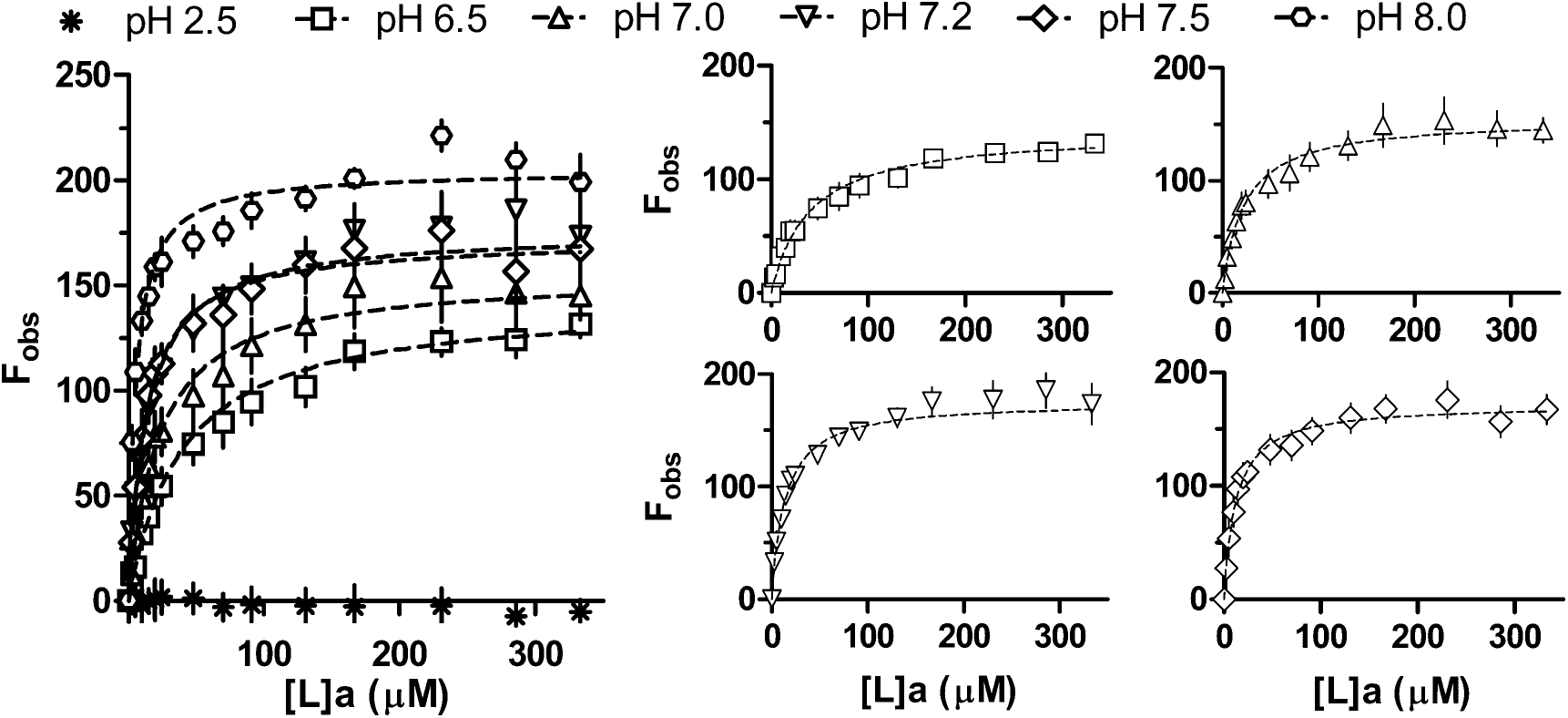
Binding curves for Ni^2+^➔*Cc*SBPII titrations (left-panel) with 10 mM HEPES at pH 2.5 (asterisk), pH 6.5 (square), pH 7.0 (upward triangle), pH 7.2 (downward triangle), pH 7.5 (diamond), and pH 8 (hexagon). Right panels show the same curves in the left panel individually (except pH 2.5 and pH 8.0) for easier visual inspection.

### microITFQ-LTA for ligand screening with *Cc*SBPII in a defined buffer

High-throughput ligand screens using thermal shift assays for SBPs have been conducted for metabolic intermediates, carbohydrates, amino acids, and sulfur compounds.^44–46^ In these screens, SBPs were tested for binding to myriad compounds. In contrast, Ni-BPs are often assumed to be isofunctional for a single metal ion and their promiscuity with other metal ions rarely ever reported. In our analysis of the peptide/nickel binding protein family (InterPro ID: IPR0306078), *Cc*SBPII clustered in the same phylogenetic branch as *Cj*NikZ and nearby other characterized Ni-BPs (data not shown). Based on this information and the structural superimposition of *Cc*SBPII to *Cj*NikZ (Supplementary Figure 2), we hypothesized *Cc*SBPII’s target metal ion is Ni^2+^. In the previous section, we provided evidence for a Ni^2+^➔*Cc*SBPII binding event with low micromolar affinity in 10 mM HEPES pH 7.2. Metal SBPs with crystal structures on the Protein Data Bank (PDB) have been solved with Co^2+^, Cu^2+^, Zn^2+^, Mn^2+^, or Cd^2+^ inside the binding cavity. Therefore, we performed a small ligand screen for *Cc*SBPII’s interactions with these other metal ions in 10 mM HEPES pH 7.2 (Figure 6). When NiCl_2_ and NiSO_4_ were used to prepare the Ni^2+^ ligand solution, similar *K*_D_ values were obtained for *Cc*SBPII: 14.13 μM with a CI of 13.27 to 14.98 μM, and 12.00 μM with a CI of 11.12 to 12.88 μM, respectively. Since they are nearly identical, and because we previously used NiCl_2_, we prepared the other metal ligands using their chloride salt to be consistent.

**Figure 6.**
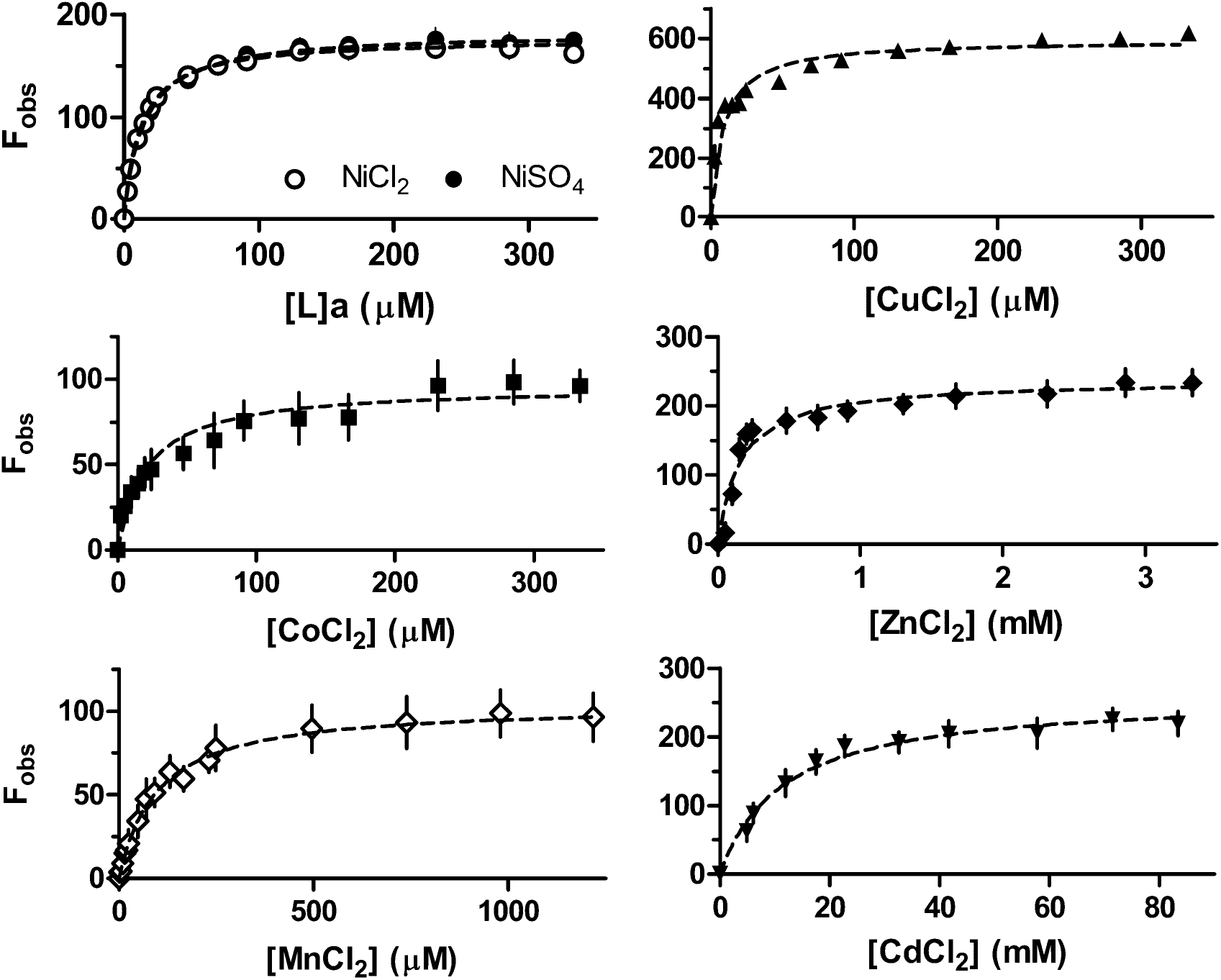
Binding curves for M^2+^➔*Cc*SBPII titrations in 10 mM HEPES pH 7.2, where M represents a divalent metal: NiCl_2_ (empty circle), NiSO_4_ (filled circle), CuCl_2_ (upward triangle), CoCl_2_ (square), ZnCl_2_ (filled diamond), MnCl_2_ (empty diamond), CdCl_2_ (downward triangle). Metal ligands are prepared in 10 mM HEPES pH 7.2 Note the differences in range for the y-axis and x-axis when comparing curves.

The results are summarized in Table 2. Binding affinities (*K*_D_) varied over 3 orders of magnitude, and the observed order of weakest to strongest affinity was Cd^2+^ ≪ Zn^2+^ < Mn^2+^ < Co^2+^ < Ni^2+^ < Cu^2+^. Notably, Cd^2+^ and Zn^2+^ are spectroscopically silent metal ions due to their complete *d* orbitals, yet they still generated a saturable signal in the microITFQ-LTA. This indicates the mechanism of fluorescence quenching was not collisional quenching, but better explained by the changes in the polarity of the environment surrounding the proteinaceous tryptophans due to binding-induced conformational changes. This is confirmed by other binding studies using specITFQ-LTA to study Cd^2+^ and Zn^2+^ interactions with metalloproteins.^47,48^

Also of note, the observed order resembles the natural order of stabilities (*i.e*. the Irving-William series).^49,50^ However, an anomaly in this observed order is Zn^2+^ appearing to be weaker in affinity than Co^2+^, whereas in the originally reported Irving-William series Zn^2+^ is predicted to have a stronger affinity than Co^2+^. The theoretical and empirical rationale for the order of the Irving-William series is based on observations made with mono-to poly-dentate amino-bearing molecules in aqueous solution, and high-spin octahedral complexes.^50^ While amino acid monomers were considered in this conceptualization, it did not consider the spatial constraints imparted by a relatively rigid protein structure that may only be able to accommodate a specific coordination geometry. Zn^2+^ is overwhelmingly found in metalloproteins with a tetrahedral geometry despite its tendency to form an octahedral [Zn(H_2_O)_6_]^2+^ complex in an aqueous solution.^51,52^ It is possible that *Cc*SBPII’s hypothesized binding residues are unable to readily position themselves to accommodate a ligand with a tetrahedral preference, perhaps requiring a conformational change to do so that would explain its 1.47-fold increase in F_max_ compared to Ni^2+^. Zn^2+^ also has a stronger preference for softer elements like sulfur in cysteine, of which is not found in the *Cc*SBPII binding cavity.

For Cd^2+^➔*Cc*SBPII, it is unclear why exactly its *K*_D_ increased over 967.5-fold compared to Ni^2+^ when Zn^2+^ only increased 12.8-fold. This dramatic difference in affinity may be due to Cd^2+^ having a marked preference for aspartate and glutamate residues, which are not included in the hypothesized binding residues, but are plentiful across *Cc*SBPII’s surface.^53^ Cd^2+^ is relatively flexible with its geometric preference compared to the other metal ions tested here. In fact, Cd^2+^ has been found in numerous crystal structures of metalloproteins in *in*completely formed trigonal planar and tetrahedral complexes,^52^ so gross adventitious binding may be occurring at sites across *Cc*SBPII that may not quench tryptophan fluorescence, similar to one SBP from *Listeria monocytogenes* (PDB ID: 5JPD). In such circumstances, x-ray crystallography is a better choice than microITFQ-LTA for study numerous interactions simultaneously.

The apparent ability for *Cc*SBPII to bind Cu^2+^ 1.4-fold more tightly than the hypothesized target Ni^2+^ and 2.5-fold more tightly than Co^2+^ is a clear demonstration of an issue bacteria face: mismetallation.^54,55^ The *ana* operon (Supplementary Figure 1) from which *Cc*SBPII is encoded from also encodes for enzymes predicted to require nickel- and cobalt-based co-factors, supporting the notion that the target is Ni^2+^ and likely Co^2+^ too. To improve Ni^2+^ selectivity, Ni-BPs have been reported to use opine metallophores akin to siderophores for iron species, suggesting that *Cc*SBPII may also naturally use chelators. These opine metallophores are comprised of a histidine connected to a pyruvate or an α-ketoglutarate, and Ni-BP crystal structures have been solved with nickel-histidine complexes inside the binding cavities, so we chose to prepare 1:2 stoichiometric ligand solutions of Cu^2+^, Ni^2+^, and Co^2+^ with L-histidine to titrate into *Cc*SBPII in 10 mM HEPES pH 7.2 (Figure 7). Using Equation (**6**) with the KK method and inclusion of n, we observed substantial changes in *K*_D_, summarized in Table 2. In all cases n is near 1, suggesting a one-binding site model is appropriate. In the absence of histidine, the observed order of affinity from weakest to strongest binding was Co^2+^ < Ni^2+^ < Cu^2+^; in its presence it was Co^2+^ < Cu^2+^ < Ni^2+^. Specifically, by introducing histidine, *K*_D_ decreased 37.8-fold for Co^2+^, 236.7-fold for Cu^2+^, and a remarkable 2728.4-fold for Ni^2+^ to obtain a *K*_D_ of 4.4 nM with a CI of 1.3 to 8.8 nM. *Cc*SBPII binds 11.6-fold more tightly to a nickel-histidine complex than *Cj*NikZ, reported earlier in this study to have a *K*_D_ of 51 nM. Further testing reveals that the imidazole ring of histidine contributes the most to the decrease in *K*_D_ compared to glycine (the backbone), but the presence of both imidazole and glycine is not sufficient enough for improving the binding affinity and selectivity for Ni^2+^ (Figure 8). This indicates it is specifically the histidine molecule, not just its components, that is required to potentially counter mismetallation by Cu^2+^. However, it is unclear whether *Cc*SBPII is binding to a nickel-histidine complex that has a 1:1 stoichiometry like that found in *Cj*NikZ, or 1:2 like those found in *YpYntA* or *Sa*NikA.^28^

**Figure 7.**
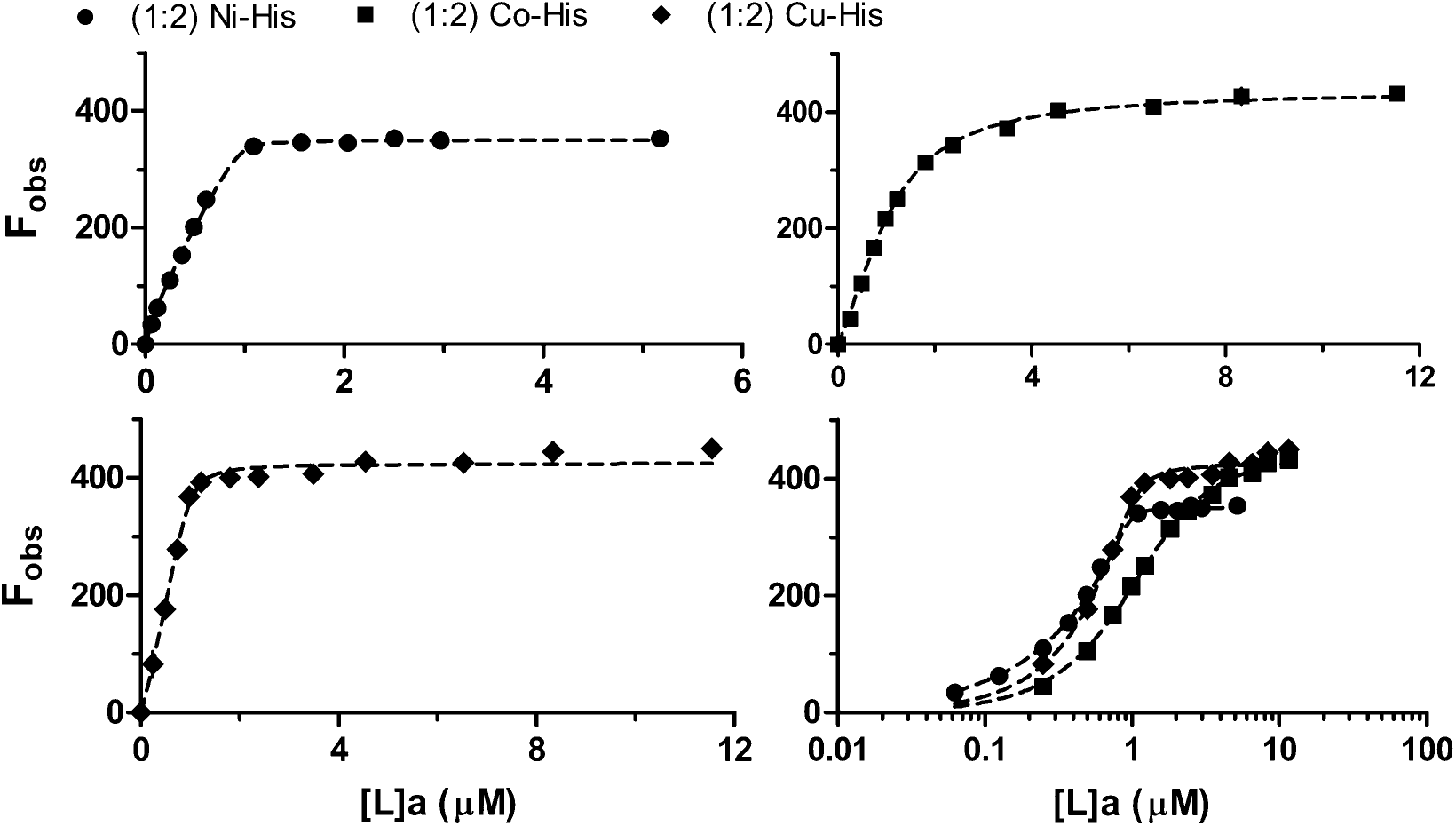
Binding curves for MHC➔*Cc*SBPII titrations in 10 mM HEPES pH 7.2, where MHC represents a metal-histidine complex prepared with a 1:2 stoichiometric ratio: nickel-histidine or Ni-His (circles), cobalt-histidine or Co-His (squares), and copper-histidine or Cu-His (diamonds). To compare the three, a plot with a logarithmic x-axis is provided for comparison (bottom-right panel).

**Figure 8.**
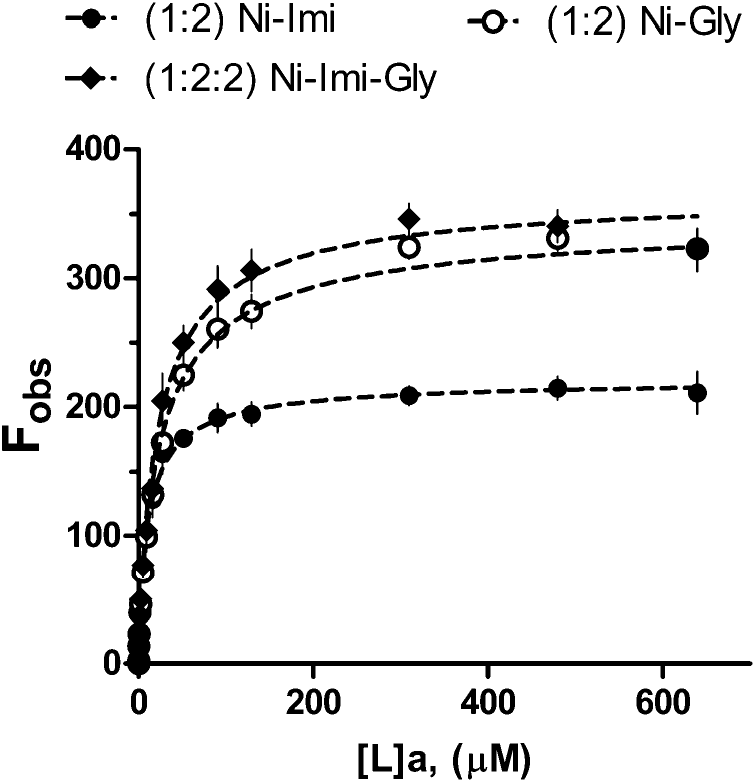
Binding curves for C➔*Cc*SBPII titrations in 10 mM HEPES pH 7.2, where C represents a nickel in complex with imidazole (1:2, Ni-Imi, filled circles), glycine (1:2, Ni-Gly, empty circles), and both (1:2:2, Ni-Imi-Gly, diamond).

### Limitations of the method

The microITFQ-LTA has several limitations that ought to be considered during experimental design and data analysis. First, the user must estimate the magnitude of [*L*]_a_ that will lead to maximal protein fluorescence quenching and prepare their ligand solution accordingly. If the user does not reach saturation in the titration range tested, best-fit parameter values will be poorly determined and lead to incorrect conclusions about the protein’s binding profile. We recommend a titration range over 3 orders of magnitude with a plateau reached in the largest order of magnitude; or alternatively, when the *F*_obs_ value ceases to increase with differences within 1 standard deviation for the last three data points. For example, in Figure 4, we did not observe saturation of *Cc*SBPII by Ni^2+^ in the presence of 100 and 1000 mM NaCl. The corresponding best-fit parameter values therefore need to be interpreted with caution. Additionally, the beginning of the binding curve has a larger effect on the *K*_D_ value, so it is important for users to have enough titration steps to accurately fit the model.

Second, Cu^2+^ is known to undergo a redox reaction with the hydroxyl group of HEPES to form Cu^+^.^56^ We attempted to measure the IFE this may cause by titrating Cu^2+^ (same preparation and concentrations used in the Cu^2+^➔*Cc*SBPII assay) into 10 mM HEPES pH 7.2 without protein and did not observe anything different from titrating the buffer alone (data not shown). Cu^+^ prefers tetrahedral geometries and lower coordination numbers,^57^ and our observation that *Cc*SBPII has a low affinity for Zn^2+^, which has a preferred tetrahedral geometry in protein structures, suggests that it is unlikely *Cc*SBPII would specifically bind to Cu^+^ as the major copper species with a *K*_D_ value (8.54 μM) less than that of Ni^2+^ (12.00 μM). Additionally, Cu^+^ requires stabilization through interactions with alkenes and amines. This means its concentrations can be very low (*viz*. its presence is negligible) without catalysts or additional chemicals to drive the redox reaction forward and maintain its presence.^58^ In our purification and activity buffers, we did not add such components. Stabilization of Cu^+^ by adventitious binding to the protein’s amine groups could occur, especially for micromolar and nanomolar *K*_D_ interactions as observed here,^59^ but this phenomenon is poorly studied and does not nullify the usefulness of well documented methods for estimating *K*_D_ values. Nevertheless, we caution future users of the microITFQ-LTA to carefully analyze data involving Cu^2+^, especially when additional buffering components are introduced that may stabilize the Cu^+^ species. To better characterize the redox state of copper bound to *Cc*SBPII, electron paramagnetic resonance (EPR) spectroscopy and nuclear magnetic resonance (NMR) are recommended.^60^

Third, the models presented here in Equations (**6**) and (**7**) are not appropriate for definitively elucidating mismetallation, and therefore directly studying the competition between metal ions. Our study of histidine’s effect on *Cc*SBPII’s ability to bind to Ni^2+^ compared to Cu^2+^ (Figure 7) must be further explored using appropriate models.^59,61^ Similarly, studying the binding stoichiometry for metal complexes (*i.e*. histidine with Ni^2+^, Co^2+^ Cu^2+^) cannot simply use the Hill Coefficient n appended to Equations (**6**) and (**7**) to definitively elucidate the ratio of protein to metal to complexing agent. We mostly observed n ≈ 1 meaning that a one-site binding model could be true in many cases, but further research is needed using other binding models,^59^ or ITC and x-ray crystallography^42^ to support these findings.

Lastly, we note that the position of tryptophan residues in the protein structure should be inspected to determine if they are solvent exposed and protruding into the metal-binding site as this could lead to collisional quenching in addition to the intramolecular self-quenching underpinning the microITFQ-LTA described here. Such a scenario would lead to a measured fluorescence quenching that would be a combination of the two phenomena, thus overestimating the association phenomenon. Structural data for proteins can be used to inspect the spatial location of the tryptophan residues computationally or experimentally.^62,63^ Correcting for collisional quenching is described elsewhere.^9^ For *Cj*NikZ and *Cc*SBPII, our structural analysis (data not shown) revealed their tryptophan residues are buried in the hydrophobic core.

## Conclusion

The validity of the microIPFQ-LTA method proposed in this study was confirmed by using it to determine the *K*_D_ value for Ni^2+^➔*Cj*NikZ and (1:2) Ni-His➔*Cj*NikZ binding events. We were able to closely replicate existing *K*_D_ values from literature and support crystal structure data. We then tested this method on *Cc*SBPII, an uncharacterized putative Ni-BP from *C. carboxidivorans*, to demonstrate how a series of microITFQ-LTAs could be designed to study the effect of buffer composition on Ni^2+^➔*Cc*SBPII binding and how *Cc*SBPII interacts with various metal ions and complexes.

Approximately 240 replicates were conducted using the microITFQ-LTA in this study. The time required to process a single 96-well plate (*i.e*. one set of titrations manually by multichannel pipette) was 1.5 hrs. 240 replicates would have required >350 hrs (at least two weeks *straight*) if performed on a one-cuvette-per-run spectrofluorometer; ITC would have taken substantially longer. While dedicated spectrofluorometers and ITC can provide more in-depth and more comprehensive analyses of binding events, high-throughput methods like the microITFQ-LTA are crucial for exploring protein libraries with large buffer and ligand screens.

## Abbreviations

LB: Luria-Bertani
TB: terrific broth
ACN: acetonitrile
TFA: trifluoracetic acid
HEPES: (4-(2-hydroxyethyl)-1-piperazineethanesulfonic acid)
IPTG: isopropyl β-D-1-thiogalactopyranosid
TCEP: tris(2-carboxyethyl)phosphine
MWCO: molecular weight cut-off
NTA: nitriloacetic acid
SDS-PAGE: sodium dodecyl sulfate polyacrylamide gel electrophoresis
TEV: tobacco endomosaic virus
FPLC: fast performance liquid chromatography
IMAC: immobilized metal-affinity chromatography
aEX: anion exchange chromatography
ITFQ-LTA: intrinsic tryptophan fluorescence quenching ligand titration assay
ICP-MS: inductively coiled plasma mass spectrometry
MALDI-ToF: matrix assisted laser desorptionionization time-of-flight
CD: circular dichroism
PDB: Protein Data Bank
IFE: Inner Filter Effect
NLLS: non-linear least squares
ITC: isothermal titration calorimetry

## Acknowledgements

The authors thank (1) Anna Khusnutdinova, Robert Flick, and Chao Chen for advice on experimental design and data processing, (2) Dr. Bridget A. Bergquist and Joan de Vera for access to and training on their Thermo-Fisher X-Series 2 ICP-MS in the Department of Earth Sciences at the University of Toronto, and (3) Dr. Walid Houry, Vaibhav Bhandari, and Elisa Leung for access to and training on the Bruker MALDI-ToF Biotyper and the JASCO J-800 spectropolarimeter in the Department of Biochemistry at the University of Toronto. This work was supported by the Ontario Ministry of Economic Development, Job Creation and Trade through the Elements of Bio-mining ORF-RE program. PD is a recipient of an Ontario Graduate Scholarship.

## Supplementary Materials

## Appendix A Additional Protocols

### Bioinformatics of the ana operon and CcSBPII

The *ana* operon of *C. carboxidivorans* was discovered through a blastp search using the *Cj*NikZ open-reading frame (UniProt ID: A0A0M4UJ91), and can be found here: tiny.cc/ccanaoperon. The locations of the promoter, ribosome binding sites, and transcriptional terminator were predicted using BPROM^64^, RBS Calculator^65^, and FindTerm^64^, respectively. The cobalamin riboswitch was predicted using Riboswitch Scanner.^66^ Multiple sequence alignments were made and visualized on JalView using the MUSCLE algorithm with default settings.^67^ The 3D structure of *Cc*SBPII was made using SWISS-MODEL with unliganded *Cj*NikZ as the template (PDB ID: 4OET), and visualized on PyMOL.^28,63^

### Matrix-assisted laser desorption/ionization time-of-flight (MALDI-ToF)

Matrix solution (30% v/v ACN, 0.05% v/v TFA, sinapinic acid to saturation) was briefly centrifuged to remove undissolved sinapanic acid, then combined with 1 μL of 1 mg/mL *Cc*SBPII in a 1:1 ratio and applied to a polished steel target plate (Bruker, Part No. 8281817) to dry at room temperature. The Bruker Microflex LT MALDI-ToF was used in the 20 – 100 kDa range to analyze the ACN:TFA-*Cc*SBPII precipitate by collecting 200 shots with the laser at 85% power and 60 Hz. Data was analyzed on flexAnalysis v3.4.

### Inductively-coiled plasma mass spectrometry (ICP-MS)

A multi-element standard curve (Millipore Sigma, #1094800100) was prepared (0, 0.01, 0.1, 0.5, 1, 5, 10, 15, and 20 ppb) volumetrically in MilliQ water, each containing a final concentration of 2% TraceMetal™ Grade HNO_3_ (ThermoFisher, #A509P500) and 8 ppb TraceCERT^®^ germanium (Millipore Sigma, #05419) for the internal standard. *Cc*SBPII samples for metal analysis were prepared in the same manner with a final *Cc*SBPII concentration of 0.67 μM (total volume = 3 mL per *Cc*SBPII sample). For metal analysis, the ICP-MS (Thermo-Fisher X-Series 2) was equilibrated at 20 °C for 30 min. The system was then flushed with 2% TraceMetal™ Grade HNO_3_ for 30 min and tuned using the tune A solution (Fisher Scientific, #NC9063415). Triplicates were acquired and processed automatically by PlasmaLab, then visualized using GraphPad Prism 5.0.

### Circular dichroism (CD)

The JASCO-800 spectropolarimeter was flushed with nitrogen gas (20 L/min) for 15 min and allowed to equilibrate at 25 °C with running distilled water before turning the lamp on. A 1-mm quartz cuvette (Millipore Sigma, #Z800015) was washed with MilliQ water, 70% EtOH, 70% HNO_3_, and a final wash with MilliQ water. 1 μM of purified *Cc*SBPII in MilliQ water was added to the cuvette and analyzed in triplicate using the Measurement Mode between 180 – 260 nm at a 1 nm interval. The curve was then smoothed using a Savitzky-Golay filter on CAPITO^68^, then visualized using GraphPad Prism 5.0.

---

**Supplementary Figure 1.**
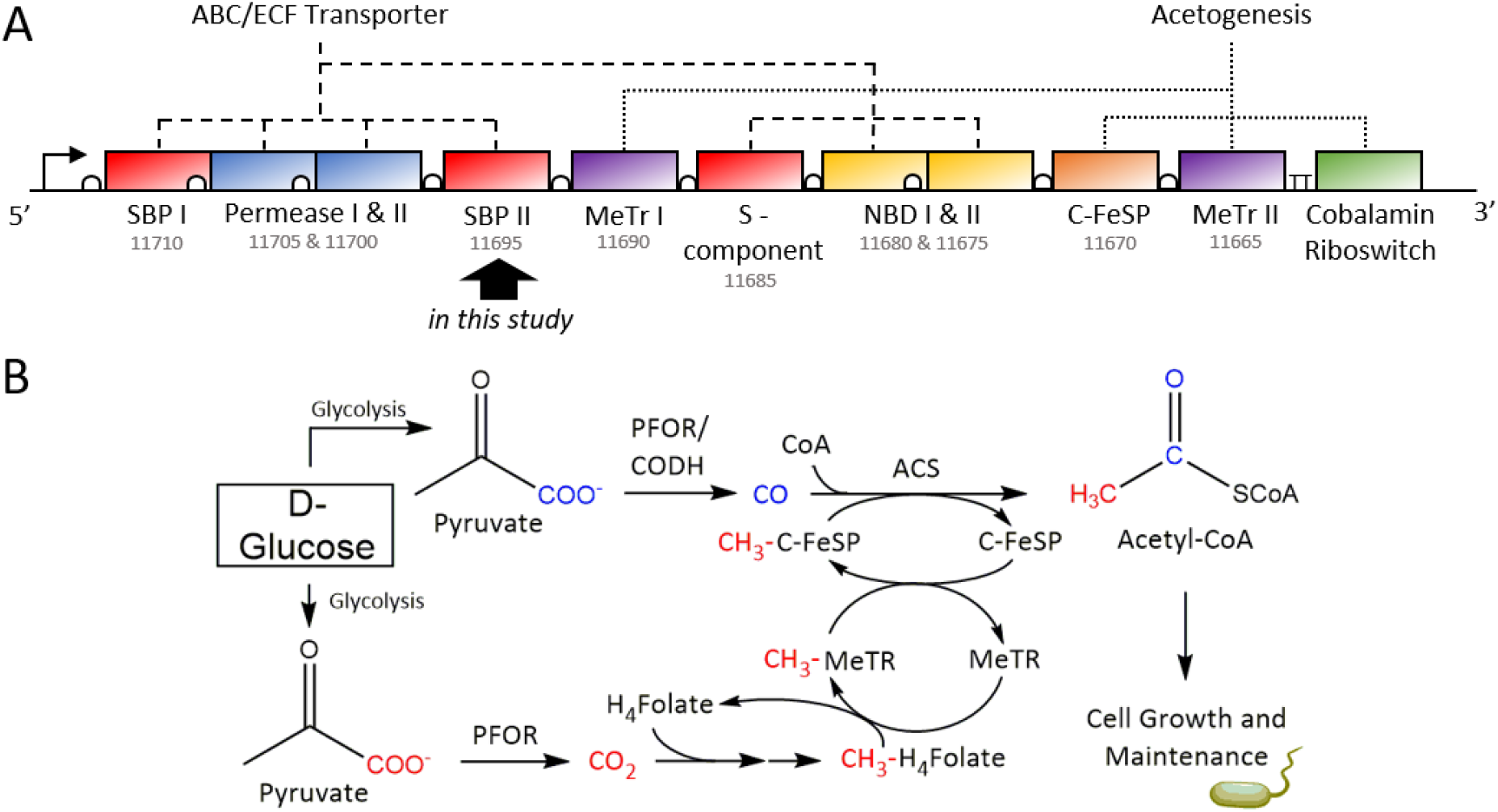
Schematic diagram of the *ana* operon from *Clostridium carboxidivorans* encoding a potential acetogenesis pathway. <u>(A)</u> The *ana* operon is comprised of two sets of coding sequences. The first encodes an ABC transporter, which includes two solute binding proteins (SBP I and SBP II), two permease units (Permease I and Permease II), two nucleotide-binding domains (NBD I and NBD II), and an energy-coupling factor S-component. The second encodes acetogenesis-related enzymes, which includes two methyltransferases (MeTR I and MeTR II), one C-FeSP, and a cobalamin riboswitch after the transcriptional terminator (TT). A promoter (right angle arrow) and ribosome binding sites (white semi-circles) indicate where the operon and coding sequences start, respectively. Numbers below coding sequence names are identifiers that can be prefixed with “Ccar_” to locate them in the *C. carboxidivorans* genome (NZ_CP011803.1). SBP II (*Cc*SBPII) is the protein of interest in this study. <u>(B)</u> Pyruvate is largely derived from glycolysis and possesses a carboxylate group (COO^-^) noted in blue and red, representing the Western and Eastern branches of the Wood-Ljungdahl pathway, respectively. Numerous enzymes require metal co-factors. PFOR requires a [4Fe-4S] cluster. CODH requires a [3Fe-4S] cluster bridged to a binuclear NiFe cluster. ACS requires a [4Fe-4S] cluster bridged to a binuclear NiNi cluster. C-FeSP requires cobalamin.

**Supplementary Figure 2.**
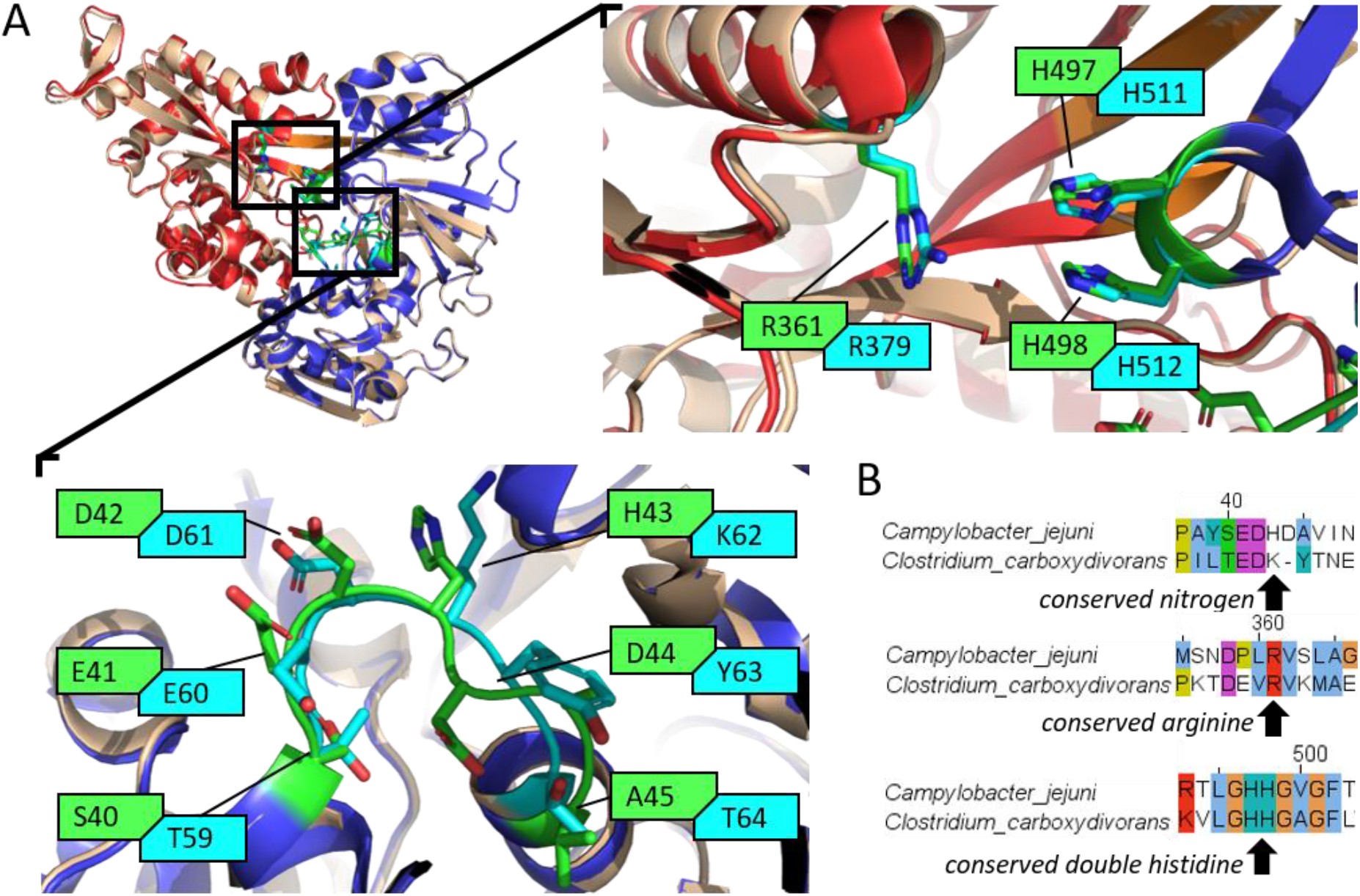
Similarities between *Cj*NikZ and *Cc*SBPII revealed by structural and sequence analysis. <u>(A)</u> A superimposition was made on PyMOL using the unliganded *Cj*NikZ structure (tan, PDB ID: 4OET) and the predicted *Cc*SBPII structure (red - Western lobe, blue - Eastern lobe, orange - hinge region) from SWISS-MODEL with 4OET as the template. The top-right panel (RHH cavity) and bottom-left panel (variable loop) are zoom-ins of the superimposition and highlight residue similarities and differences between *Cj*NikZ (green residues) and *Cc*SBPII (cyan residues). <u>(B)</u> A multiple sequence alignment reveals *Cc*SBPII uses K62 instead of H43 from *Cj*NikZ, but has conserved binding residues otherwise (black arrows).

**Supplementary Figure 3.**
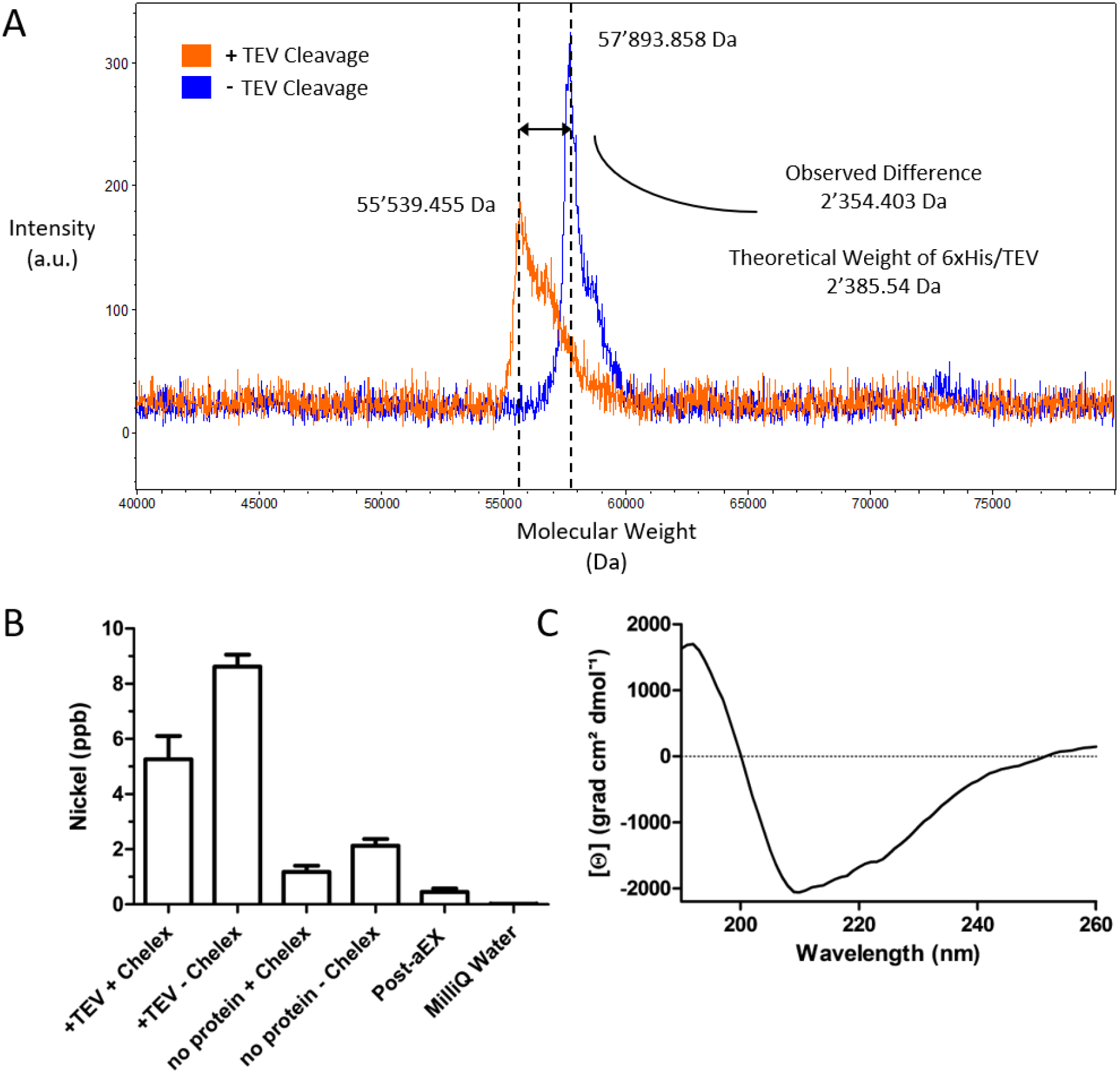
Post-purification validation of tag removal, nickel removal, and proper protein folding of purified *Cc*SBPII. <u>(A)</u> MALDI-ToF analysis of protein molecular weight with two samples (orange – addition of TEV, blue – no addition of TEV). Dashed black lines depict the maximum peaks for each sample. <u>(B)</u> ICP-MS analysis of nickel concentration to evaluate the usage of Chelex 100 to remove heavy metals from the Co-IMAC protein eluent, which is collected after applying the cell lysate to the Co-NTA and washing the resin with washing buffer. This protein eluent is combined with TEV for cleavage and dialyzed for 72 hrs with and without Chelex 100, labelled “+TEV +Chelex” and “+TEV -Chelex”, respectively. A negative control is included where the cell lysate volume is replaced with binding buffer to determine background nickel contamination by the buffer. No TEV is added to the eluent in this case and is dialyzed in the same fashion, labelled “no protein + Chelex” and “no protein – Chelex”. *Cc*SBPII after the final 24 hr dialysis, prior to freezing, was also evaluated by ICP-MS and labelled “Post-aEX”. (C) CD spectrum of *Cc*SBPII after being blanked with 10 mM HEPES, pH 7.2.

**Supplementary Table 2.**
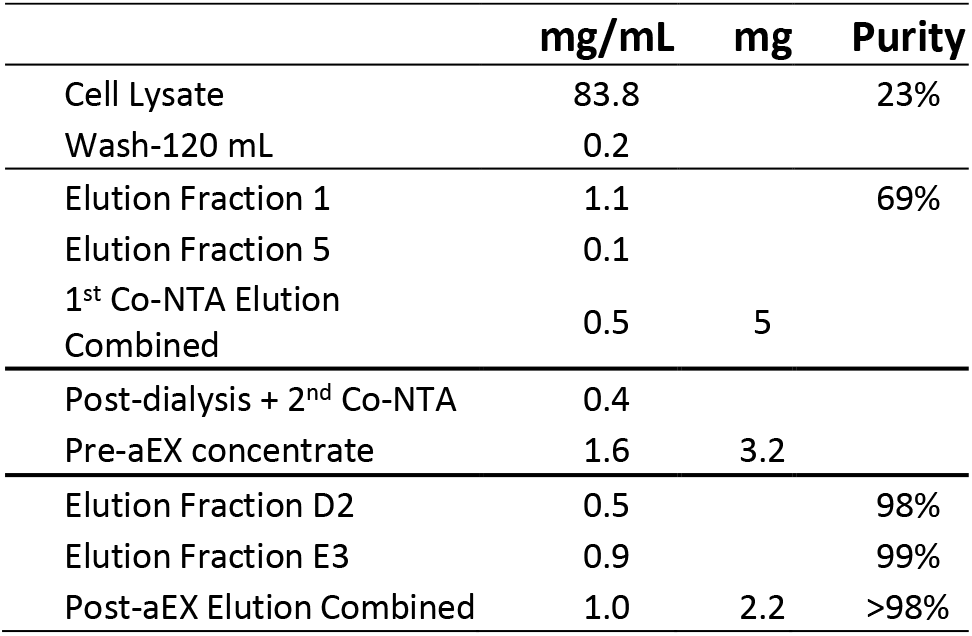
*Cc*SBPII purification and purity profile

**Supplementary Figure 4.**
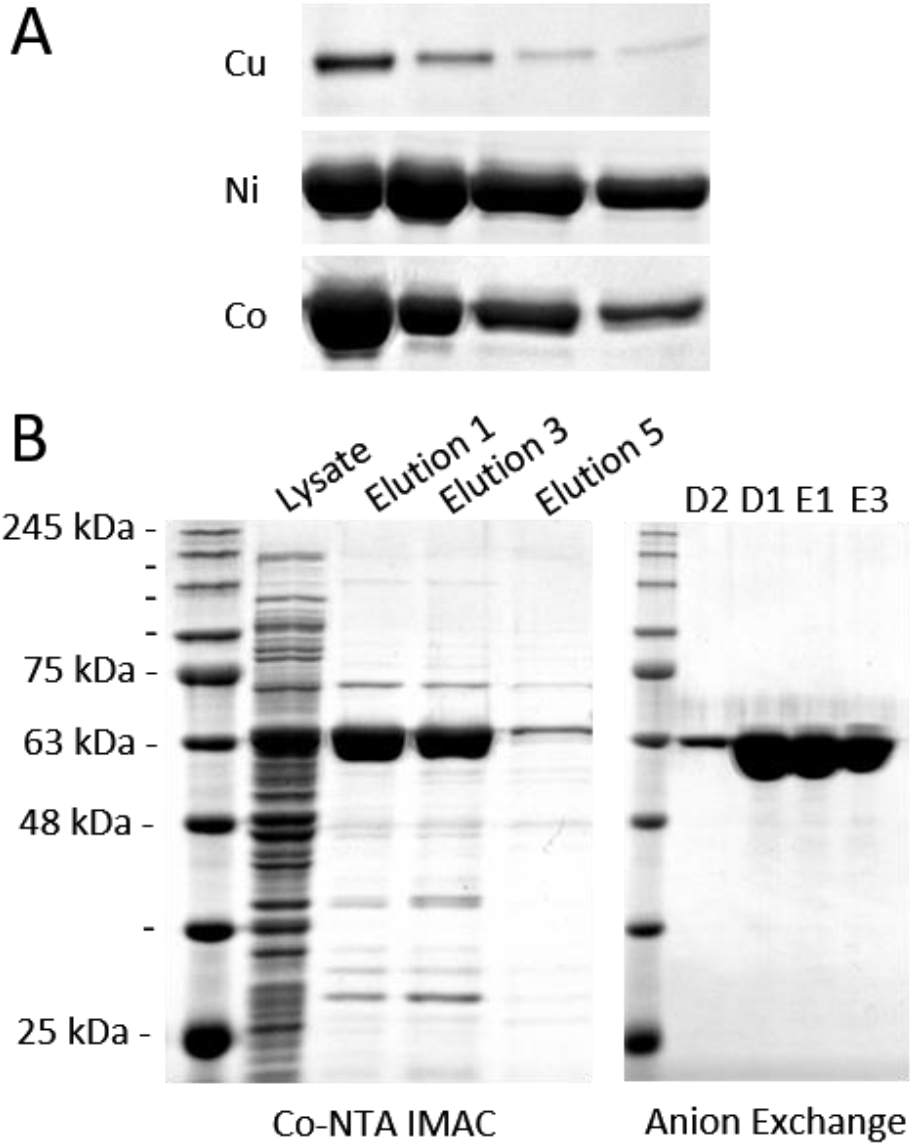
SDS-PAGE analysis of *Cc*SBPII purification. <u>(A)</u> A comparison of *Cc*SBPII purification using NTA resin charged with copper, nickel, and cobalt. <u>(B)</u> Two-step purification of *Cc*SBPII. Left-panel: 1^st^ step, purification using Co-NTA resin (elution with 250 mM imidazole). Right-panel: 2^nd^ step, purification using anion exchange chromatography (aEX) on MonoQ to remove remaining impurities.

**Supplementary Figure 5.**
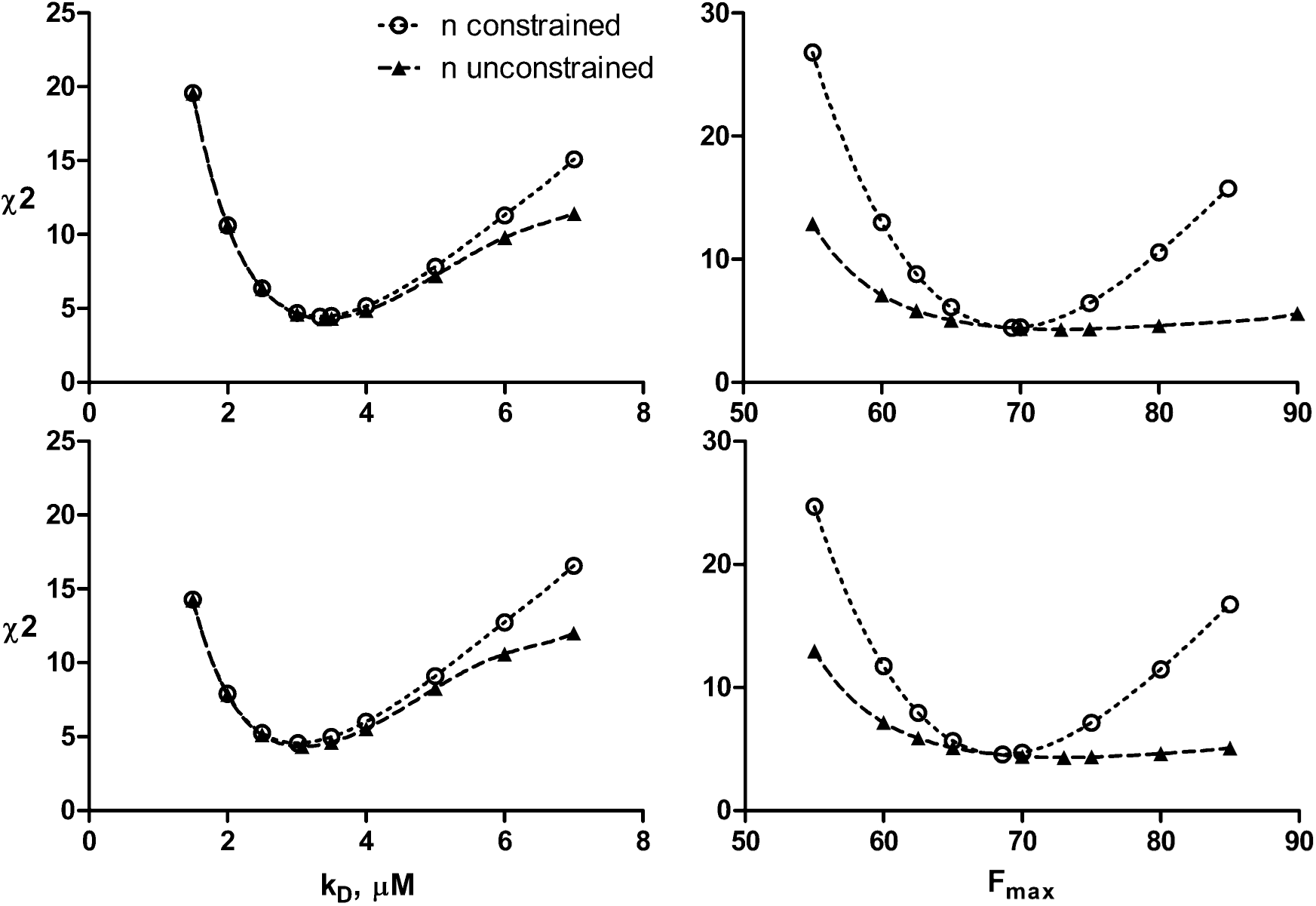
χ^2^ plots of Ni^2+^➔*Cj*NikZ to compare how n constrained (n = 1, circles) and n unconstrained (triangles) affects the behavior of global minima for best-fit *K*_D_ and *F*_max_ parameters. Top-Left Panel: χ^2^ vs. *K*_D_ using Equation (7). Top-Right Panel: χ^2^ vs. *F*_max_ using Equation (7). Bottom-Left Panel: χ^2^ vs. *K*_D_ using Equation (6). Bottom-Right Panel: χ^2^ vs. *F*_max_ using Equation (6). Curves of best-fit for these plots are polynomial functions generated from Excel, and solved using Wolfram Alpha to generate CIs. These plots are created following instructions described by the KK method. Graph generated on Prism.

**Supplementary Figure 6.**
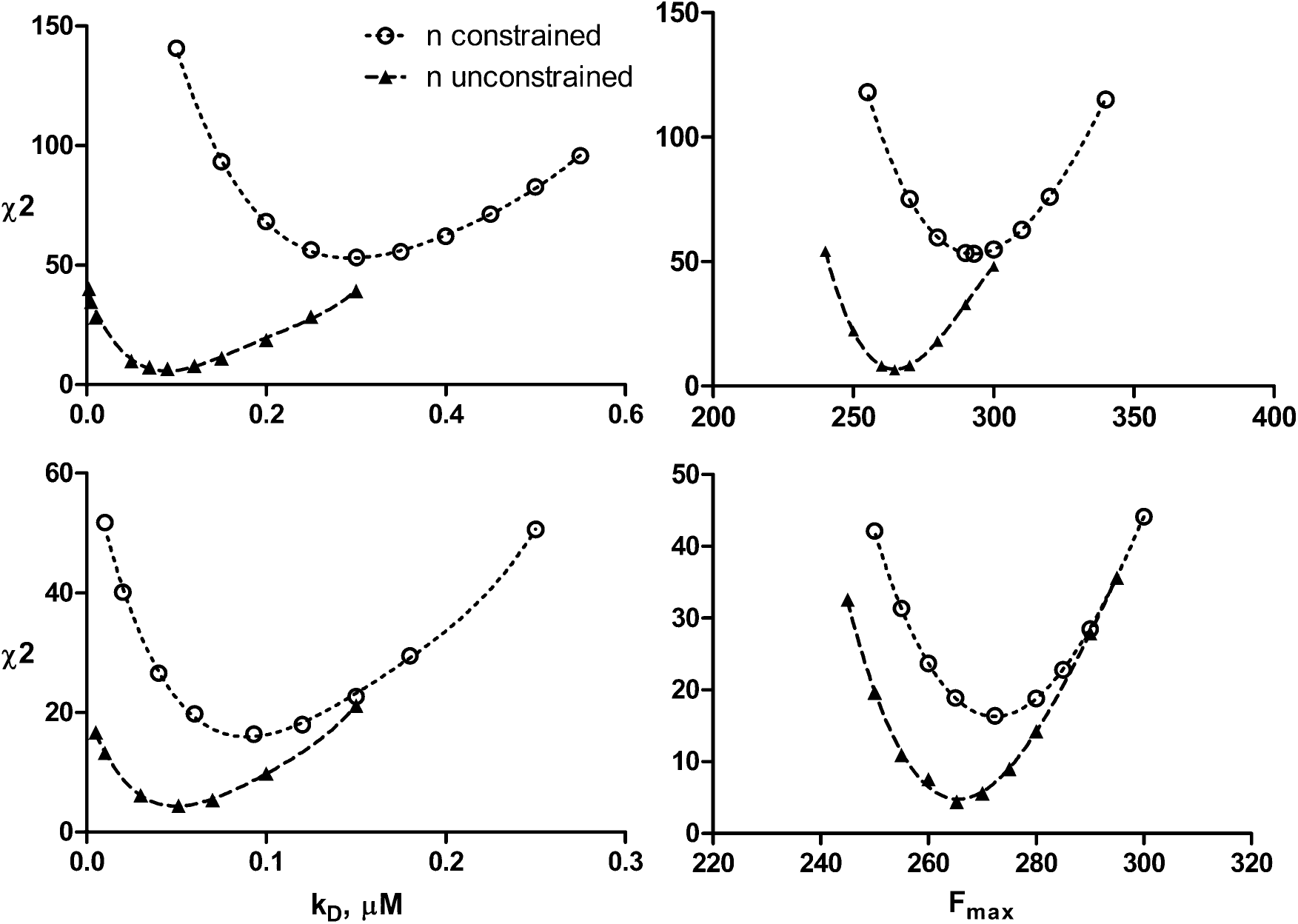
χ^2^ plots of (1:2) Ni-His➔*Cj*NikZ to compare how n constrained (n = 1, circles) and n unconstrained (triangles) affects the behavior of global minima for best-fit *K*_D_ and *F*_max_ parameters. Top-Left Panel: χ^2^ vs. *K*_D_ using Equation (7). Top-Right Panel: χ^2^ vs. *F*_max_ using Equation (7). Bottom-Left Panel: χ^2^ vs. *K*_D_ using Equation (6). Bottom-Right Panel: χ^2^ vs. *F*_max_ using Equation (6). Curves of best-fit for these plots are polynomial functions generated from Excel, and solved using Wolfram Alpha to generate CIs. These plots are created following instructions described by the KK method. Graph generated on Prism.

**Supplementary Figure 7.**
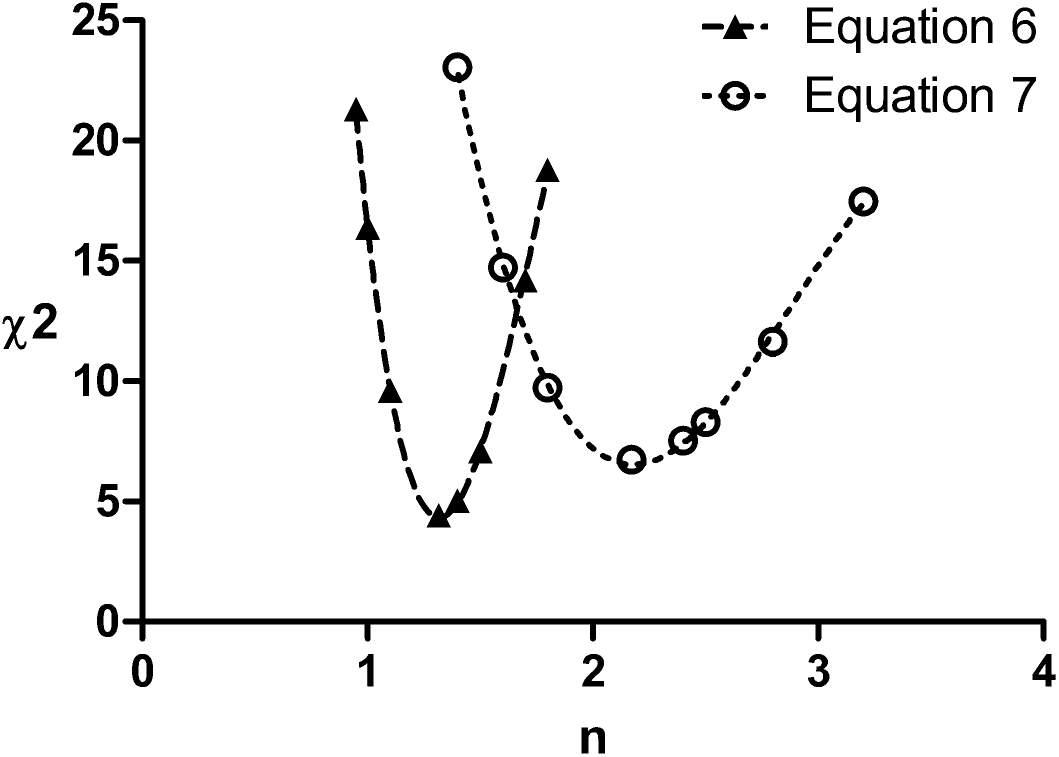
χ^2^ plot of (1:2) Ni-His➔*Cj*NikZ to compare how the inclusion of n in Equations (6) and (7) alters the global minimum for the parameter n. Curves of best-fit for these plots are polynomial functions generated from Excel, and solved using Wolfram Alpha to generate CIs. These plots are created following instructions described by the KK method. Graph generated on Prism.

**Supplementary Figure 8.**
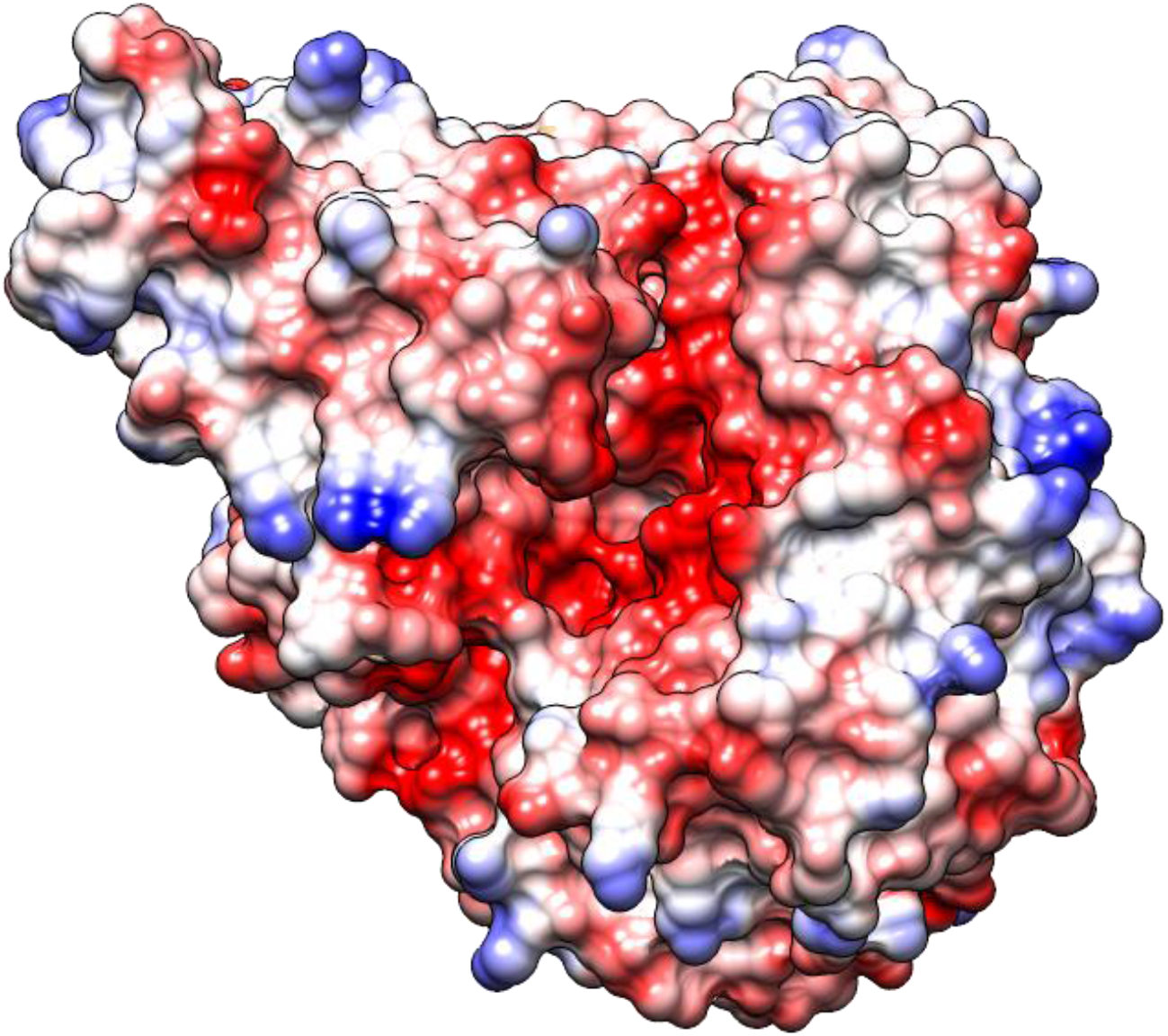
Coulombic surface view of *Cc*SBPII (homology model from *Cj*NikZ, PDB ID: 4OET) highlighting regions of positive charge (blue), negative charge (red), and neutral (white). Visualized on Chimera 1.13.1.

